# Altered drug metabolism and increased susceptibility to fatty liver disease in myotonic dystrophy

**DOI:** 10.1101/2021.04.06.438688

**Authors:** Zac Dewald, Andrew Gupta, Ullas V. Chembazhi, Auinash Kalsotra

**Affiliations:** Department of Biochemistry, University of Illinois, Urbana-Champaign, IL, United States; Cancer Center@Illinois, University of Illinois, Urbana-Champaign, IL, United States; Carl R. Woese Institute for Genomic Biology, University of Illinois, Urbana-Champaign, IL, United States

## Abstract

Myotonic Dystrophy type 1 (DM1), a prevalent muscular dystrophy affecting 1 in 2800 individuals, is associated with a toxic (CTG)_n_ repeat expansion in the *DMPK* gene. While DM1 affects multiple systems, recent studies highlight its link to liver pathology, glucose intolerance, and drug sensitivity. Our study focused on liver implications by creating a hepatocyte-specific DM1 mouse model. Expression of toxic RNA in hepatocytes sequestered muscleblind-like (MBNL) proteins, impacting hepatocellular activity. DM1-induced liver alterations included morphological changes, inflammation, necrosis, and fatty accumulation. Impaired drug metabolism and clearance were evident in DM1 mice and increased susceptibility to diet-induced fatty liver disease. Notably, alternative splicing of acetyl-CoA carboxylase 1 induced excessive lipid accumulation in DM1 livers, exacerbated by high-fat, high-sugar diets. These findings unveil disruptions in hepatic functions, predisposing DM1 livers to injury, fatty liver disease, and compromised drug clearance. Understanding these mechanisms is crucial for addressing the complex health challenges in DM1 patients and optimizing treatment strategies.

## Introduction

Myotonic Dystrophy Type 1 (DM1) is an autosomal dominant disease and the second most common form of muscular dystrophy, affecting more than one in three thousand adults in North America^1–3^. The cardinal symptoms of DM1 include myotonia, debilitating muscle weakness and wasting, abnormal heart function, and excessive fatigue^1,2^. Despite DM1’s initial characterization as a form of muscular dystrophy, the disease is genuinely multisystemic; patients report various gastrointestinal, metabolic, and neurological dysfunctions, such as excessive daytime sleepiness and insulin resistance^2,4^.

DM1 is caused by a (CTG)n repeat expansion in the 3’ UTR of a ubiquitously expressed gene Dystrophia Myotonica protein kinase (*DMPK*)^5–7^. The (CUG)n containing RNAs resulting from the transcription of the diseased *DMPK* gene form hairpin secondary structures and aggregate in the nucleus, forming discrete RNA foci^7,8^. These foci interact with and sequester the muscle blind-like (MBNL) family of splicing factors^9,10^. MBNL proteins affect many developmentally-regulated alternative splicing and polyadenylation decisions in various tissues throughout the process of maturation towards adulthood; thus, their loss-of-activity in DM1 shifts splicing of target pre-mRNAs towards preadolescent-like patterns, inducing specific features of the disease^11–14^. This reversal of transcriptomic patterns from mature-to-immature state drives many disease symptoms to become more prevalent later in life, with diagnosis often occurring in the mid to late thirties^12,15,16^. However, diagnosis typically occurs only after the development of significant muscular or neurological symptoms, which allows for the subtle and long-term consequences of the disease to go unmanaged^16^.

DM1 patients are abnormally sensitive to a wide range of anesthetics and muscle relaxants, resulting in prolonged anesthesia recovery, heightened pulmonary dysfunction, and, in some cases, death^1,17,18^. The disruption of neurological and muscular function, hallmarks of DM1, is often blamed for this sensitivity. However, within the last twenty years, several studies have demonstrated that DM1 patients have an increased susceptibility to non-alcoholic fatty liver disease (NAFLD), metabolic syndrome, and liver damage^19–22^. These studies would suggest inappropriate liver function and a predisposition for liver injury in DM1 patients. A malfunctioning liver could also help explain the sensitivity to anesthetic treatment; a liver that cannot provide adequate metabolism of xenobiotic material would prolong the clearance time for many drugs and may increase their potency. Minor malfunctions in liver response to metabolic signaling and drug metabolism may frequently occur in DM1 patients; however, none have investigated this possibility.

Here, we sought to determine what effects DM1 might have on hepatic functions and overall liver health. Utilizing two previously established mouse lines, we generated a mouse model in which we induced the expression of (CUG)960 repeat-containing RNA, specifically in the hepatocytes within the liver^23–25^. Combining acute and long-term DM1 liver mice models with systematic biochemical, molecular, and high-resolution transcriptome analyses, we found that toxic (CUG)960 RNA expression triggered global gene expression and RNA processing defects in the hepatocytes. These transcriptome defects led to various physiological and cellular pathologies, including accumulation of lipids and fatty liver, increased susceptibility to insult and injury, and misregulation of xenobiotic metabolism. Specifically, we uncovered that aberrant splicing and upregulation of Acetyl CoA Carboxylase 1 (ACC1), the rate-limiting enzyme for *de novo* fatty acid biosynthesis, drives the NAFLD phenotype in DM1-afflicted mice livers. Importantly, both the fatty liver and the poor drug metabolism phenotypes are intrinsic to the repeat RNA toxicity within hepatocytes, as these defects were not detected in transgenic mice, which express the repeat RNA only in the muscle tissues. Thus, our study reveals that DM1 disrupts normal hepatic functions, predisposes the liver to fatty liver disease and injury, and confirms the need for further research into the effects of DM1 in non-traditional tissues, including the liver.

## Results

### A hepatocyte-specific murine model of DM1 recapitulates the molecular features of the disease in the liver

The pathogenic mechanism of DM1 is comprised of three primary parts: i) the transcription and production of a long CUG repeat-containing RNA, ii) the accumulation of this RNA into nuclear foci, and iii) the sequestration of MBNL proteins into such RNA foci, which results in the decrease of MBNL directed RNA processing activities^26–28^. To study the effects of DM1 within the liver, we generated a bi-transgenic murine model by combining two existing mice models. First is the tetracycline-inducible mouse model with a *DMPK* transgene containing the last five exons of human *DMPK* and 960 interrupted CTG repeats (labeled here as CUG960i RNA), developed by Cooper and colleagues^23^. The second model utilizes the expression of a reverse tetracycline trans-activator (rtTA) driven by a liver-specific apolipoprotein E (ApoE) promoter, which is highly expressed in hepatocytes^24^. By crossing these two mice models, we generated a double homozygous bi-transgenic line that allows for conditional, doxycycline (Dox)-dependent expression of the CUG960i RNA specifically in the liver tissue, thus allowing the study of DM1 disease in the liver (Fig. 1a). From now on this bi-transgenic model is referred to as the DM1 liver model. The control mice for this model contain only the homozygous ApoE-rtTA allele.

**Fig. 1.**
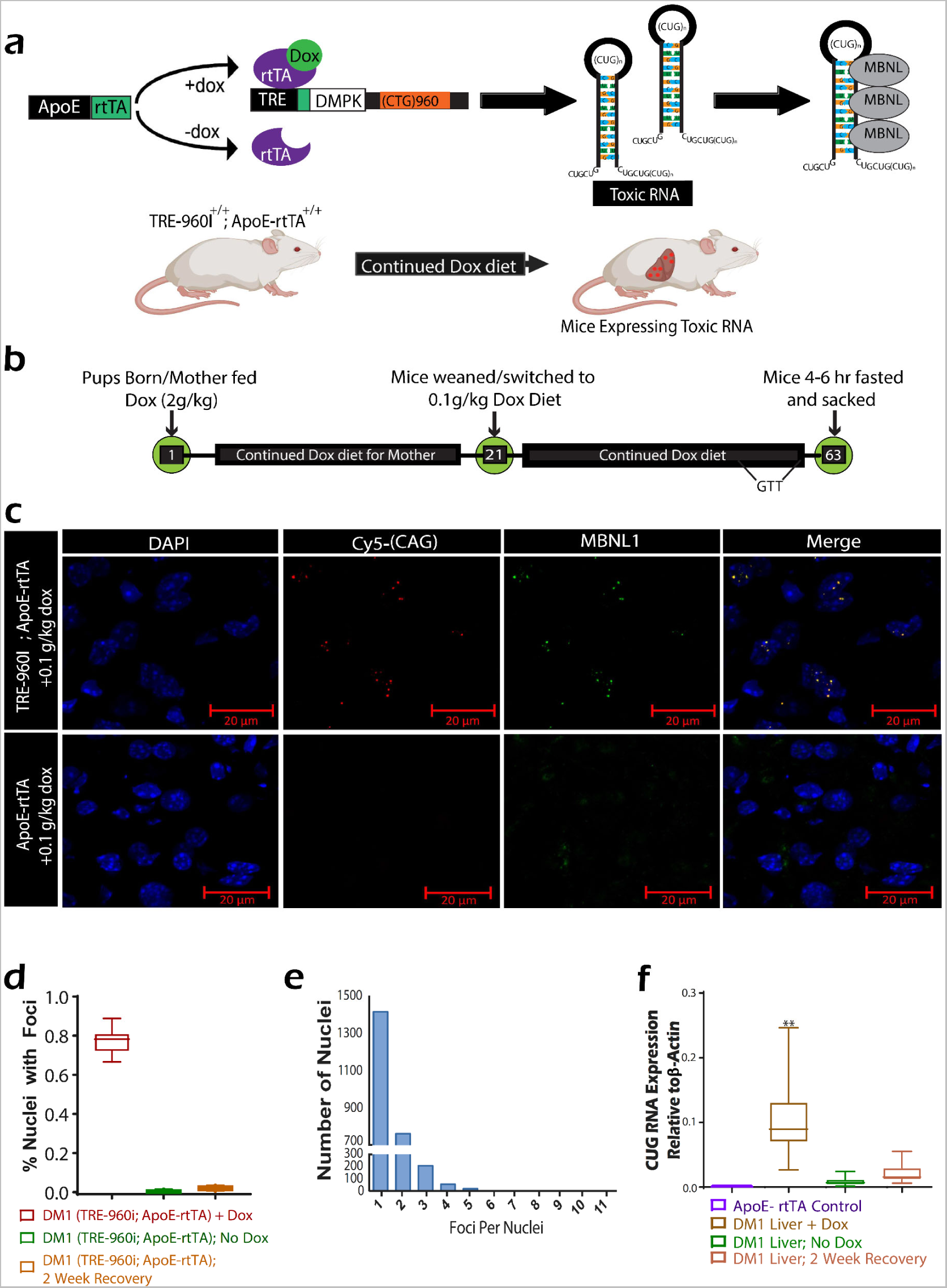
Murine Model Recapitulates Molecular Mechanism of DM1 in Hepatocytes. (a) Schematic illustrating the bi-transgenic, hepatocyte-specific, doxycycline (Dox)-inducible model developed to express toxic CUG960i RNA in mouse livers. The (CUG)_n_ repeat-containing transcripts sequester RNA-binding proteins, including MBNL proteins. Administration of Dox triggers the expression of toxic RNA in hepatocytes. This model is referred to as the DM1 liver model. (b) Experimental protocol for Dox diet feeding to induce DM1 in mouse livers, involving a 2g/kg Dox-supplemented diet until weaning on day 21, followed by a switch to a 0.1g/kg Dox diet or maintenance on the 2g/kg Dox diet for six weeks. Glucose tolerance testing (GTT) occurs a week before sacrifice. (c) Hybrid RNA fluorescent in-situ hybridization immuno-fluorescence (RNA FISH-IF) imaging depicts toxic (CUG)_n_ RNA (red) and Mbnl1 (green) foci in hepatocyte nuclei (blue). (d) Quantification of CUG960i/Mbnl1 foci in hepatocyte nuclei via RNA FISH-IF. DM1 liver mice on 0.1g/kg Dox diet (n=7) are compared to the No Dox diet (n=5) and 2-week recovery mice (n=7). (e) Distribution of CUG960i/Mbnl1 foci per hepatocyte nucleus in mice fed 0.1g/kg Dox diet for one-month post-weaning (n=7). (f) Quantitative-PCR (qPCR) analysis of CUG960i RNA in hepatocytes and whole liver of DM1 liver mice and controls (ApoE-rtTA: n=5, DM1 liver mice: n=20, No-Dox mice: n=9, 2-week recovery mice: n=7). Box plots display first to third quartile with median line; mean ± SD for others. **P < 0.01.

To mimic the DM1 conditions seen in human patients, we induced the disease in newborn pups by feeding the mother a diet supplemented with 2 g of Dox per kg of chow. Once weaned, mice then continued a Dox diet at a lower dose of 0.1 g/kg until they reached adulthood at nine weeks (Fig. 1b). At eight weeks of age, mice were fasted for 20-22 hours and administered a glucose tolerance test (GTT).

To confirm the appropriate expression of the CUG960i RNA and the formation of toxic RNA/RNA binding protein (Rbp) foci, we utilized fluorescent in-situ hybridization and immunofluorescence (FISH-IF) imaging with probes targeting the CUG repeat sequence within the RNA (Fig. 1c). Bright puncta of condensed CUG RNA were seen in the nuclei of most hepatocytes. The signal from these RNA foci overlapped with the immunofluorescent signal when fluorescent MBNL1 or MBNL2 antibodies were used, indicating that the toxic CUG RNAs have successfully sequestered the MBNL proteins. The MBNL-containing RNA foci only occurred in the DM1 mice’s livers and not in the ApoE-rtTA control mice’s livers.

Quantification of the CUG960i/MBNL foci indicated that over 80% of hepatocyte nuclei in the DM1 mice contain at least one RNA focus, ensuring the uniformly distributed expression of both transgenes (Fig. 1d). The appearance of RNA foci was Dox-dependent — when mice with both the ApoE-rtTA and CUG960i alleles were not fed Dox or if the Dox diet was withdrawn for a week or more, the RNA foci were undetectable. The distribution of CUG960i RNA foci per nuclei followed a Poison curve, with most hepatocytes having one to three foci and a small number of hepatocytes exceeding ten plus foci within a single nucleus (Fig. 1e).

Finally, to assay the amount of toxic RNA produced, CUG960i transgene expression was quantified by isolating RNA from whole livers and conducting qPCR using primers located in the final exon of the *Dmpk* transcript. DM1 liver mice expressed the CUG960i transgene at levels near ten percent of *β-actin* transcripts within the liver, compared to ApoE-rtTA control mice, which showed no evidence of CUG960i expression (Fig. 1f). A comparison of CUG960i RNA within isolated hepatocytes versus that of the whole liver confirmed that the transgene’s expression occurs primarily within hepatocytes (Supplementary Fig. 1a, b). Livers of the bi-transgenic mice not fed Dox showed no CUG960i expression, confirming that Dox must be provided to these animals to express sufficient amounts of toxic RNA. (Fig. 1f).

### Expression of CUG960i RNA induces global transcriptomic changes within hepatocytes

Upon establishing that the DM1 liver model reproduces the molecular features of the disease, we prepared total RNA from purified hepatocytes isolated from the ApoE-rtTA controls and DM1-afflicted mice fed a 2.0 g/kg Dox-supplemented diet for nine weeks. We next assessed the splicing patterns of MBNL1-regulated exons within these RNA samples using end-point reverse-transcription PCR (RT-PCR) assays. The DM1 mouse livers consistently reproduced an alternative splicing pattern that significantly deviated from the control samples (Fig. 2a), confirming that, like the muscle and brain tissues, the expanded CUG repeat-containing RNA of DM1 also induces splicing defects in the liver.

**Fig. 2.**
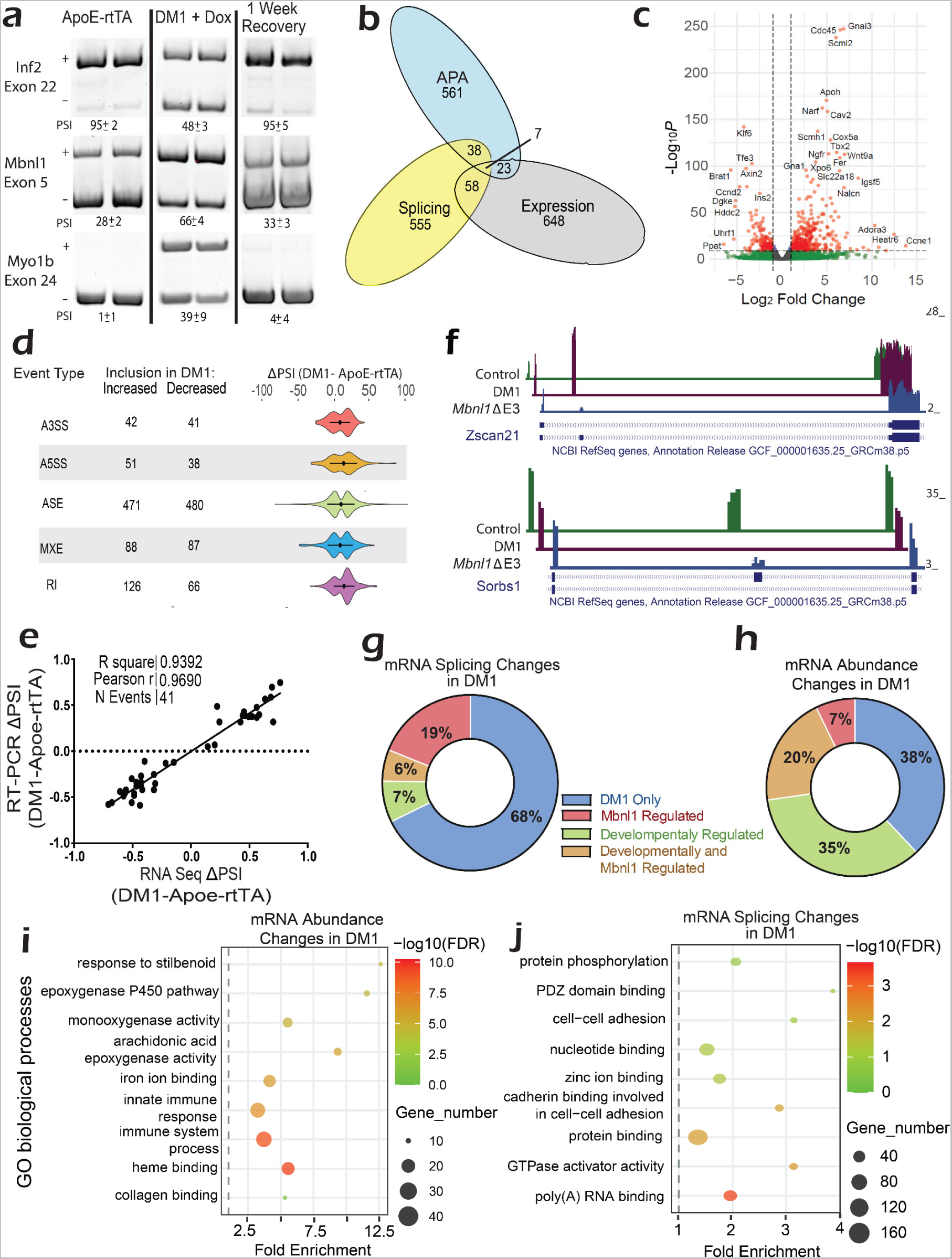
DM1 Induces Global Transcriptomic Alterations in Hepatocytes. (a) RT-PCR splicing analysis of select MBNL1 targets. Bands indicate exon presence or absence, with (+) for inclusion and (-) for exclusion. Targets are listed to the left of the image, with the percentage “spliced in” (PSI) below. (b) Overlap of alternative splicing, alternative polyadenylation (APA), and expression changes upon DM1 induction in hepatocytes. (c) Volcano Plot illustrating mRNA abundance changes from RNA-seq (d) Violin plots displaying inclusion levels of alternative splicing events from RNA-seq data. MXE = mutually exclusive, A3SS/ A5SS = alternative 3’/5’, ASE = alternative cassette exon, and RI = retained intron. (e) Change in PSI determined by RT-PCR (y-axis) vs. RNA Seq analysis (x-axis) for 30 events. (f) Gene Tracks of representative genes showing alternative exon inclusion in DM1 liver, *Mbnl1*^ΔE3/ΔE3^ (KO), or wild-type animals. (g) Pie chart – comparison of alternatively spliced genes regulated by DM1, Mbnl1, or maturation in the liver. (h) Pie chart – comparison of differentially expressed genes (DEG) regulated by DM1, Mbnl1, or maturation. (i) GO Diagram - mRNA Processing: Selected processes related to genes with alternative mRNA processing events in DM1 liver. (j) GO Diagram - Differential Expression: Processes related to genes undergoing differential expression in DM1 liver. n=3 for DM1 liver and controls, n=2 for MBNL1KO and controls.

We next performed high-resolution RNA sequencing of poly(A) selected RNA from the purified hepatocyte samples to further explore the genome-wide RNA processing defects in DM1-afflicted livers. Analysis of the resulting data revealed widespread changes in the DM1 hepatocyte transcriptome, with significant changes in mRNA abundance, splicing, and alternative polyadenylation (ApA) (Fig. 2b). Focusing on gene expression, inducing DM1 within the murine liver changed the mRNA abundance of 760 transcripts at a 2-fold level or higher, with 516 upregulated and 244 downregulated compared to control livers (Fig. 2c).

As the MBNL proteins are most known for regulating alternative splicing events, it is not surprising that nearly one thousand splicing events changed upon the expression of the CUG960i RNA within the liver. Of the 928 splicing events that demonstrated a greater than 10% change in PSI (Percent Spliced In) within the DM1 liver, every type of alternative splicing event was represented, with most falling under the category of cassette exons, and 35 of those events showing a ΔPSI change of 50% or higher (Fig. 2d, Supplementary Fig. 1c). Using RT-PCR, forty-one of these alternatively spliced events were validated; the comparison showed a high consistency between the RNA-seq and RT-PCR results (Fig. 2, e).

Gene ontology analysis of the transcripts experiencing dysregulation in abundance, splicing, or ApA revealed enrichments in unique functional categories. Transcripts changing in abundance were enriched in glucose, lipid, and energy-related metabolism, as well as oxo-reductase and cytochrome p450 activities (Fig. 2i, Supplementary Table 1)^29^. The transcripts with altered splicing patterns were enriched in mRNA processing, signal transduction, and protein phosphorylation. A substantial number of transcripts encoding proteins associated with the immune response, specifically the response to viral and bacterial infection, exhibited defects in both overall abundance and splicing (Fig. 2j, Supplementary Tables 1-3). Transcripts with misregulated ApA events, much like misregulated splicing events, were enriched in nucleotide binding, protein binding, and transport-related functions.

The proposed molecular mechanism of DM1 entails disrupting MBNL protein activities, resulting in a transcriptomic shift away from the normal state of healthy adult tissue and towards an immature state in the muscles, heart, and neurons^13,27,28,30,31^. To test whether this pattern holds within the liver, we isolated hepatocytes from *Mbnl1*^ΔE3/ΔE3^ (*Mbnl1* knockout) mice and corresponding littermate wild-type controls at ten weeks of age^32^. Again, RT-PCR splice assays were performed on the poly(A) selected RNAs purified from wildtype and *Mbnl1* KO hepatocyte samples, and the results were compared to the DM1 liver and ApoE control samples. Notably, the DM1 liver samples showed a shift in splicing away from controls in the same direction as the *Mbnl1* KO samples (Fig. 2f; Supplementary Fig. 1d, e). However, the DM1 samples often demonstrated a more significant deviation from “normal” than the *Mbnl1* KO samples.

This prompted a full comparison of differentially expressed genes and alternatively spliced events between the DM1 and *Mbnl1* KO hepatocyte transcriptomes via RNA-Seq. Of the 705 alternatively spliced events in DM1 hepatocytes and the 508 that occurred in the hepatocytes of *Mbnl1* KO mice (difference in ΔPSI > 15%), only 175 events were common to both sets (Supplementary Fig. 1f). A similar pattern was observed for differentially expressed genes, with only a 15% overlap between the DM1 and *Mbnl1* KO data sets (Supplementary Fig. 1g). This limited overlap was surprising, as MBNL1 sequestration is a crucial driver of the transcriptomic defects in DM1 heart and muscle tissues.

As MBNL proteins promote tissue maturation and function, we next compared the transcriptomic changes in hepatocytes isolated from adult livers of DM1, *Mbnl1* KOs, and wild-type mice livers during postnatal maturation^33^. By comparing alternatively spliced transcripts that change in either the context of DM1, *Mbnl1* KO, or liver maturation, we found that only about 25% of the events changing in DM1 were regulated by MBNL1 (Fig. 2g, h). Of note, whereas only a modest portion of mis-spliced events in DM1 were developmentally regulated, over 50% of transcripts changing in abundance in the DM1 liver were also developmentally regulated.

To explore the limited overlap between the hepatic transcriptomes of DM1 and *Mbnl1* KO mice with developing livers, potential compensatory mechanisms for MBNL1 function were investigated. We found that while MBNL1 levels are wholly depleted within the *Mbnl1* KO livers, MBNL2 levels were elevated 4-fold, as confirmed by western blot analysis (Supplementary Fig. 1h). The upregulation of MBNL2 might explain why there is lower-than-expected overlap between changes in the *Mbnl1* KO and DM1 liver models. Additionally, the increase of MBNL2 implies a compensatory mechanism that buffers the effects of Mbnl1 loss within the liver, a mechanism demonstrated in other tissues^34,35^.

### Hepatocyte-specific expression of CUG960i RNA induces increased lipid accumulation and liver injury

As the effects of DM1 in the liver are unstudied, and even the role of MBNL proteins in the liver is unknown, we took a generalized approach to assess the pathological consequences of DM1 within the liver. This process started before sacrifice, as blood glucose levels just before sacrifice indicate a slight difference in the blood glucose levels between male DM1 liver mice and male controls (Fig. 3a). However, this difference does not occur within the female groups (Supplementary Fig. 2a). There was also no difference between DM1 liver mice and controls during glucose tolerance testing performed in the weeks before sacrifice (Fig. 3b; Supplementary Fig. 2b). Median mouse weight between female control and DM1 mice also showed no significant difference; however, DM1 male mice were 10-17% larger than ApoE-rtTA control males (Supplementary Fig. 2c). This increase in size is likely not due to the induction of DM1 in the DM1 liver mice, as male mice of this strain were larger than the ApoE-rtTA control males regardless of whether they were consuming a Dox diet.

**Fig. 3.**
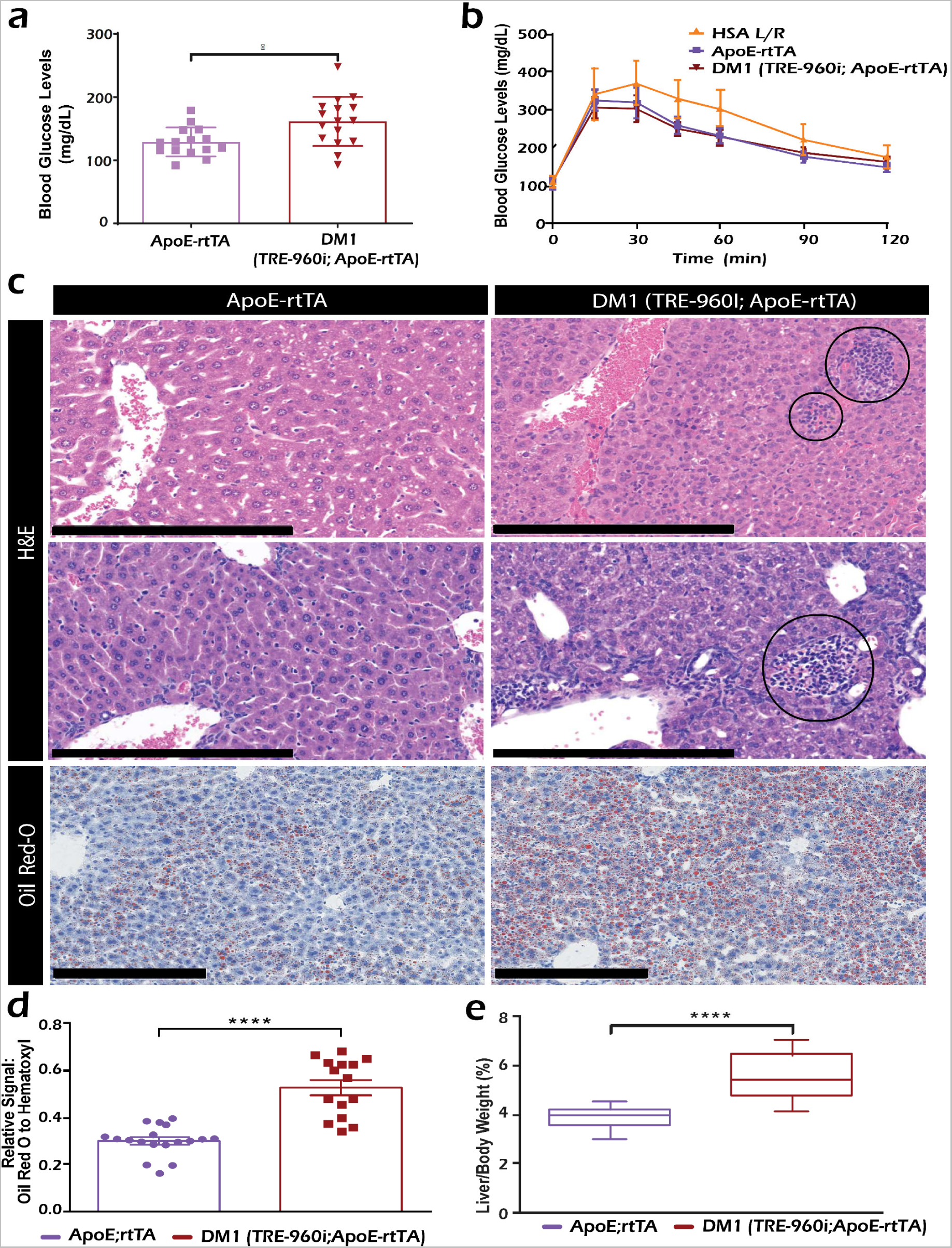
DM1 Induces Hepatic Lipid Accumulation and Injury. (a) Blood glucose levels measured before sacrifice after a 4-hour fast (n=15 for ApoE-rtTA mice and 16 for DM1 mice). (b) Glucose tolerance testing (GTT) curves depicting blood glucose levels post-intraperitoneal glucose injection. GTT was performed a week before harvest following a 24-hour fast (n=14 for ApoE-rtTA mice, 13 for DM1 liver mice, and 18 for HSA L/R mice). (c) Representative histological images of ApoE-rtTA control and DM1 mice. Hematoxylin and eosin (H&E) images showcasing inflammation and necrosis in the DM1 liver, with inflammation and necrosis circled, are at the top. Oil Red-O images indicating lipid droplets (Red) with nuclei stained in hematoxylin (blue) are at the bottom. Black scale bars represent 200 µm. (d) Hepatosomatic index for DM1 mice and ApoE-rtTA controls (n=16 for ApoE-rtTA mice and 32 for DM1 liver mice). (e) Quantification of Oil Red O signal, relative to hematoxylin-stained nuclei, indicating lipid accumulation (n=18 for ApoE-rtTA mice and 15 for DM1 liver mice). Box plots show first to third quartile with median line; mean ± SD for all others. *P < 0.05, ****P < 0.0001.

As glucose intolerance is a common symptom in DM1, we compared GTT analysis from the DM1 liver mice and control mice against the HSA L/R mice, a DM1 model commonly studied for skeletal muscle pathologies^8^. The HSA L/R model expresses the toxic CUG repeat-containing RNA only within the muscle tissues, allowing us to compare the direct contributions of liver and muscle tissue toward glucose intolerance in DM1. While the HSA L/R mice showed significant glucose intolerance, the DM1 liver mice showed normal glucose handling (Fig. 3b).

Histological analysis of the DM1 mice livers using Hematoxylin and Eosin (H&E) staining revealed varying degrees of morphological changes and regions with decreased sinusoidal spacing within the DM1 livers (Fig. 3c). Additionally, increased lobular inflammation and necrotic patches were found within the DM1 livers (Fig. 3c).

DM1 patients have shown an increased susceptibility to fatty liver disease^19,20^. Therefore, we used Oil Red O staining on frozen mice liver tissues to interrogate the lipid accumulation within the DM1 liver model. Relative to the control animals, DM1 liver mice showed a significant increase in lipid droplets (Fig. 3c, d). While a long-term Dox diet can result in a modest accumulation of lipids in the liver, the DM1 mice consistently displayed higher lipid levels, nearly twice that of respective controls. Furthermore, the mouse liver-to-carcass weight ratio showed a significantly higher hepatosomatic index in DM1 liver mice than in controls (Fig. 3e), with a median increase of 36.6%.

### DM1 liver model mice demonstrate decreased drug metabolism

An often-reported challenge when treating DM1 patients is their increased susceptibility to anesthetics and analgesics^2,17,36^. These complications are most noticeable during surgical procedures wherein DM1 patients exhibit much longer recovery times from various anesthetics and muscle relaxants. In the case of a few anesthetics, the patient may require intervention to prevent death^18,37^. Because the liver is the primary organ involved in drug metabolism, we hypothesized that DM1 livers might be compromised in responding to and metabolizing xenobiotics, thereby decreasing DM1 patients’ ability to clear certain drugs from their system.

We first chose zoxazolamine, a muscle relaxant, to test this hypothesis. Zoxazolamine testing in mice consists of inducing muscle paralysis in the animals via zoxazolamine injection and then monitoring them until they can self-right and move around freely (Fig. 4a). Zoxazolamine metabolism shows a sex-specific response in mice; however, in both males and females, DM1 mice took at least 50% longer to recover from the drug-induced paralysis (Fig. 4b).

**Fig. 4.**
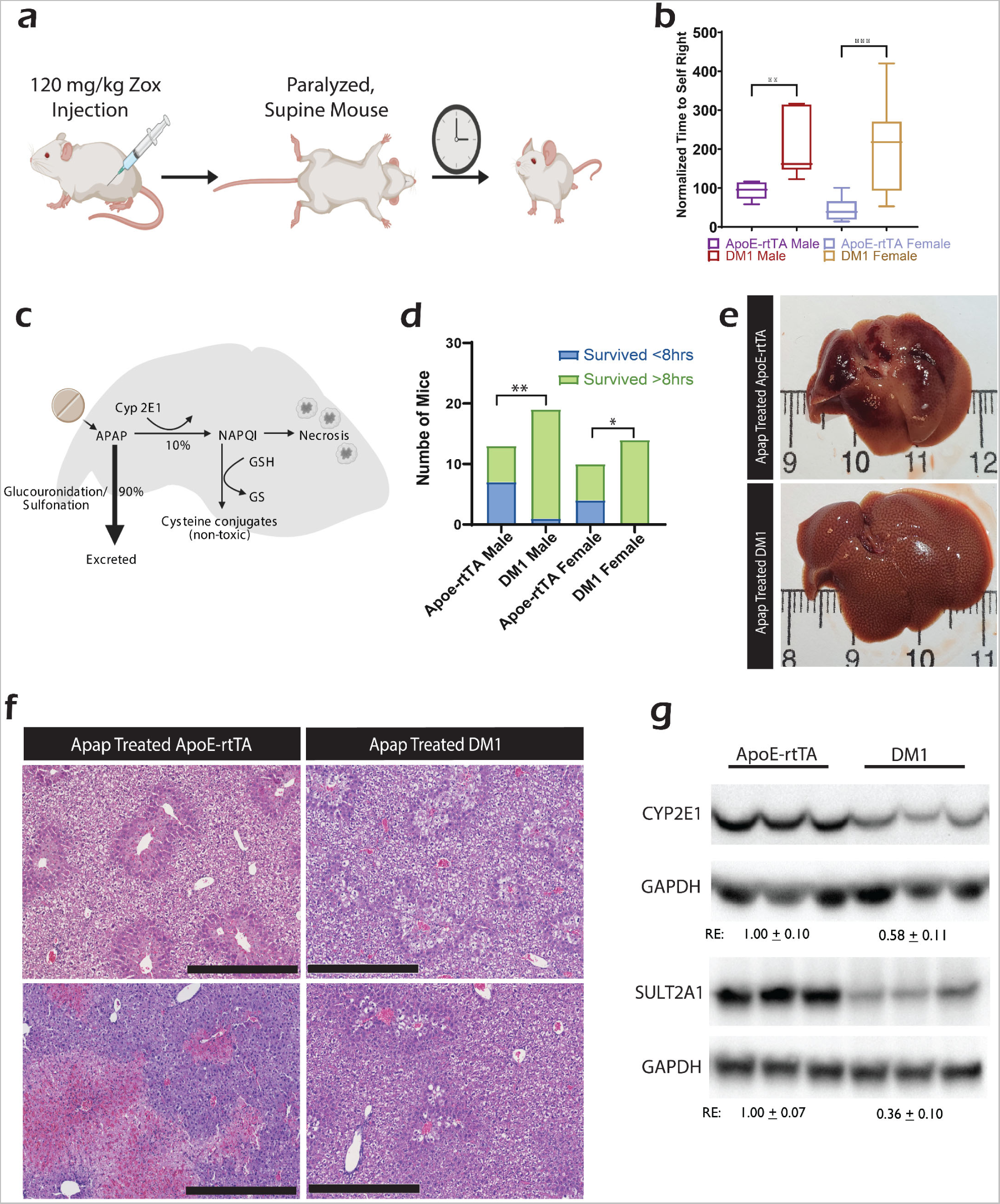
Impaired Drug Metabolism in DM1 Liver Model Mice. (a) Schematic illustrating zoxazolamine (Zox) testing procedure, measuring self-righting time post-IP injection of 120 mg/kg Zox. (b) Normalized average self-righting time of Zox-injected mice (n=9, 10, 7, 8 from left to right). (c) Diagram outlining APAP drug metabolism in hepatocytes. (d) Mortality rate 8 hours post 350 mg/kg APAP injection (n=13, 19, 10, 14 from left to right). (e) Representative livers of male mice surviving 24 hours post-APAP injection. (f) H&E images of livers from male mice 24 hours post-APAP injection. Black scale bars represent 200 µm. (g) Western blots showing CYP2E1 and SULT2A1 protein levels in control and DM1 mice. Box plots display first to third quartile with median line. *P < 0.05, **P < 0.01, ***P < 0.001.

We next tested if a similar reduction in drug metabolism occurred with common, over-the-counter analgesics such as acetaminophen (APAP). APAP causes severe liver injury/damage if consumed in high concentrations as it generates toxic levels of N-acetyl-p-benzoquinone imine (NAPQI) metabolite after oxidation by CYP2E1 in perivenous hepatocytes^38^ (Fig. 4c). Even a low dose of APAP can induce liver toxicity if CYP2E1 activity is high. Conversely, the liver can be insulated from APAP toxicity if CYP2E1 activity is ablated^39^. In most mice strains, the LD50 of APAP is between 320 and 370 mg per kg of body weight when administered intraperitoneally^40^. To see if DM1 changed susceptibility to APAP-induced hepatic injury, we injected 350 mg of APAP per kg of body weight into fasted DM1 and control mice. Mice were monitored for 8 hours and left to recover for 16 hours before being sacrificed.

A difference was immediately noticed between DM1 and control mice, as significantly more control mice died within the first 8 hours compared to DM1 mice (Fig. 4d). Upon collecting the liver from the surviving mice, 24 hours post-APAP injection, almost all control animals show widespread signs of liver necrosis (Fig. 4e). However, APAP-treated DM1 livers showed fewer instances of injury and necrosis. H&E staining of the APAP-treated livers also showed marked differences between DM1 and control mice, with DM1 mice still showing extensive injury and hepatocyte vacuolization but far less necrosis (Fig. 4f). Western blot analysis of cytochrome P450 2E1 (CYP2E1), a key enzyme involved in the metabolism APAP into NAPQI^38,41^, showed that CYP2E1 is expressed at significantly lower levels in the livers of DM1 afflicted mice (Fig. 4g). Similarly, sulfotransferase 2a1 (SULT2a1), an enzyme associated with phase 2 drug metabolism^42,43^, was also downregulated in the livers of the DM1 mice.

### DM1 murine liver models are more susceptible to fatty liver disease and injury

As DM1 patients face dietary and mobility challenges that often require counseling and careful monitoring, we set out to test if the macronutrient composition of the patient’s diet impacts the DM1 liver’s susceptibility to NAFLD^2,4,44,45^. To do so, we fed DM1 liver mice, and ApoE-rtTA controls standard chow supplemented with a 2 g/kg Dox diet until weaning, as previously described. Once weaned, the mice were switched to a high-fat, high-sugar, and heightened cholesterol (western) diet supplemented with 0.1 g/kg Dox for eight additional weeks (Fig. 5a)^46,47^. As before, we analyzed GTT and four-hour fasting glucose levels before sacrifice.

**Fig. 5.**
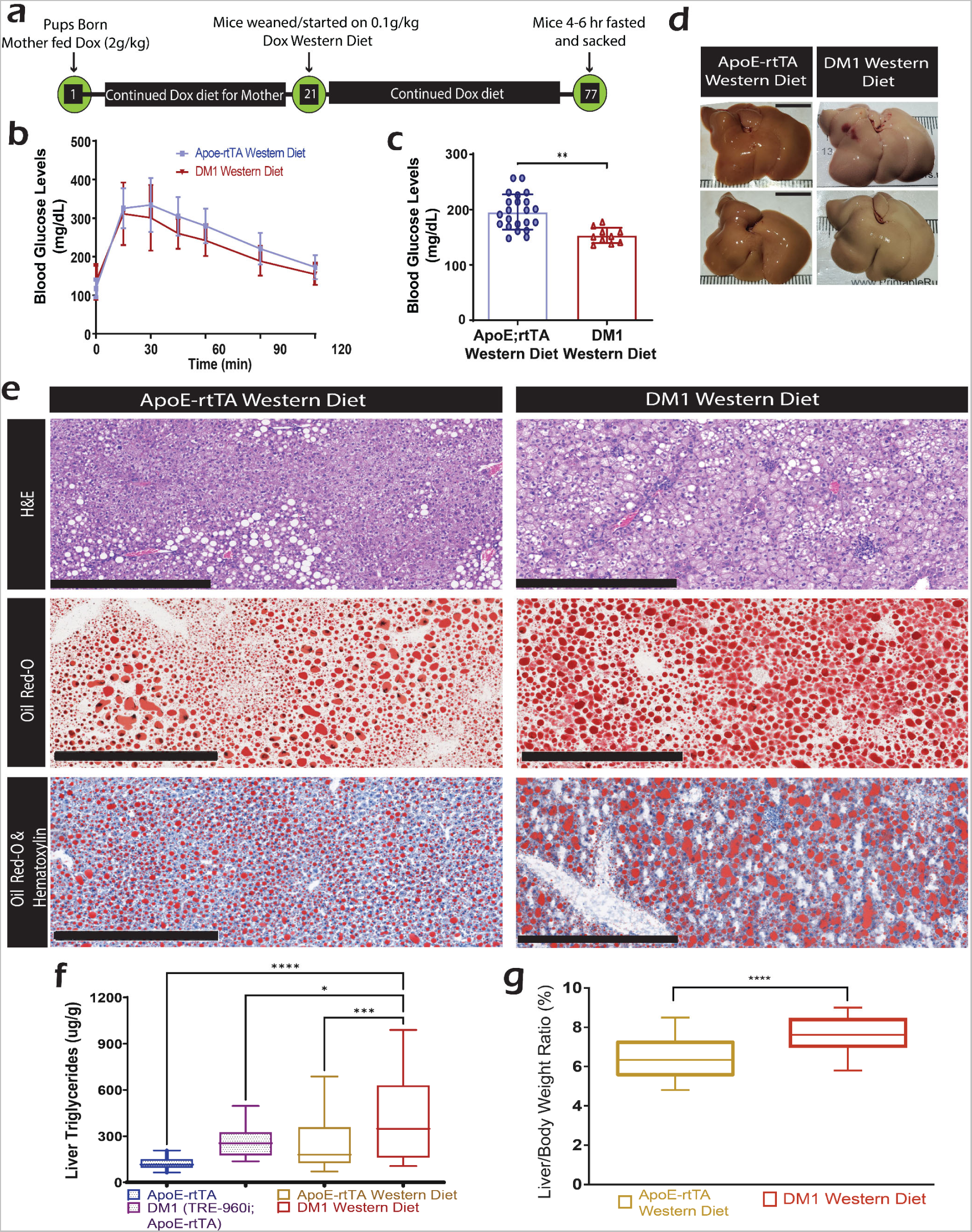
DM1 Exacerbates Diet-Induced NAFLD Severity. (a) Schematic of the feeding protocol for mice on a Western diet, starting with a 2g/kg Dox-supplemented diet at birth and transitioning to a 0.1g/kg Dox-supplemented high-fat, high-sugar “Western” diet at weaning, maintained for eight weeks until sacrifice. (b) GTT curves of male mice on the western diet, performed twice with a six-day interval (n=23 for ApoE-rtTA mice and 16 for DM1 mice). (c) Blood glucose levels before sacrifice after a 4-hour fast (n=22 for ApoE-rtTA mice and 10 for DM1 liver mice). (d) Representative liver images from ApoE-rtTA (left) and DM1 liver mice (right) on the Western diet. (e) Representative histological images, including H&E (Top) images with circled inflammation and necrosis in DM1 liver and Oil Red-O (Bottom two) images staining lipid droplets (Red) with nuclei in blue. Black scale bars represent 200 µm. (f) Analysis of extractable triglycerides in livers of mice on basal and western diets (n=21 for ApoE-rtTA mice and 14 for DM1 liver mice). (g) Hepatosomatic index for Western diet-fed DM1 mice and ApoE-rtTA mice (n=34 for ApoE-rtTA mice and 19 for DM1 liver mice). Values displayed as median to quartiles for box plots; mean ± SD for all others. *P < 0.05, **P < 0.01, ***P < 0.001.

GTT analysis again showed no difference between DM1 mice and control animals (Fig. 5b); however, there was a slight difference in 4-hour fasted blood glucose levels between male DM1 and male control mice (Fig. 5c). In reverse of the basal diet, DM1 mice had significantly lower blood glucose.

While control livers turned pale following a high fat, high sugar diet, the DM1 livers became exceedingly lighter, with much of the usual red color replaced with off-white due to excess lipid accumulation (Fig 5d). Both DM1 and control mice showed significant increases in micro-and macro-vesicular steatosis, inflammation, and evidence of cell death on the western diet compared to the regular chow diet (Fig. 5e). However, DM1 mice showed more macro-vesicular steatosis and patchy necrosis as well as ballooning and feathery degeneration after western diet feeding.

Oil Red O staining showed that livers of DM1 mice had a much higher density of lipid droplets and a significant increase in the number of large lipid droplets (Fig. 5e), making them challenging to quantify via image analysis. Therefore, we used an alternative method to determine the relative accumulation of lipids in the western diet-fed control and DM1 mice. A hexane/isopropanol lipid extraction protocol collected hydrophobic fatty acids from small liver portions. A colorimetric assay for triglycerides was performed on these extracts (Fig. 5f). This analysis revealed two features. First, the livers from DM1 mice fed the regular rodent diet accumulated as many triglycerides as those from control mice on the western diet. Second, the livers of DM1 mice fed a western diet accumulated significantly more triglycerides than any other group.

Western diet-fed DM1 mice also had a more significant increase in the liver-to-body weight ratios than controls (Fig. 5g). The mean body weight between DM1 and control mice is invariant, suggesting that the increased accumulation of lipids in the liver is not due to a more significant increase in body weight (Supplementary Fig. 2e).

### Upregulation of an alternatively spliced ACC1 isoform drives lipid accumulation in DM1 liver

To investigate how DM1 affects liver-specific lipid handling/metabolism and leads to a fatty liver, we focused on DM1-related changes in splicing/abundance of transcripts linked to lipogenesis, lipid transport, lipid metabolism, and NAFLD. Of these, acetyl-CoA carboxylase 1 (*Acc1*) particularly stood out (Fig. 6a). ACC1 is at the rate-limiting step for the conversion of excess citrate into free fatty acids (Fig. 6b); thus, any changes in its function or regulation could directly lead to excess lipid production and potentially explain the steatosis noted in the DM1-afflicted livers. ACC1 regulation involves multiple phosphorylation clusters and a necessary dimerization event to function^48–53^. Upon activation, ACC1 converts acetyl-CoA to malonyl-CoA, a necessary building block for fatty acid synthesis and lipogenesis. ACC1 activity is negatively regulated through phosphorylation by the kinases AMPK, PKA, and CDK and by excess palmitoyl-CoA levels. In contrast, it is positively regulated by insulin-induced dephosphorylation via the phosphatase PP2A and excess citrate levels (Fig. 6b)^54–56^.

**Fig. 6.**
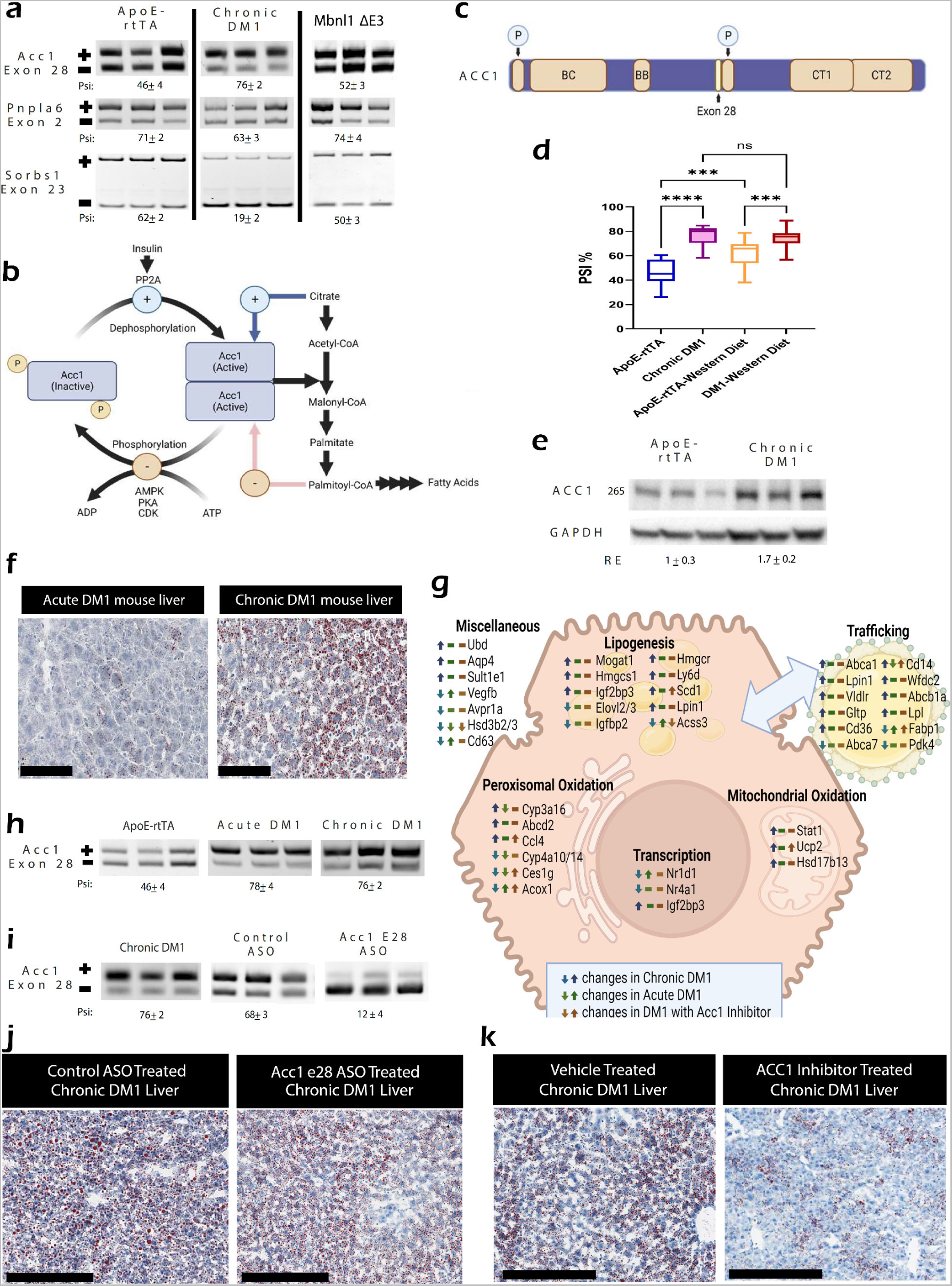
DM1 Disrupts Liver Lipid Regulation and Drives Lipogenesis via Acetyl-CoA Carboxylase 1. (a) RT-PCR splicing gel analysis of select alternative splicing events in DM1-afflicted livers. Target names are listed on the left, with PSI values below. (b) Diagram of acetyl-CoA carboxylase 1 (ACC1) regulation and function. ACC1, phosphorylated by kinases like AMPK and PKA, is inactivated. Upon insulin-driven PP2A dephosphorylation, ACC1 becomes active, converting acetyl-CoA into malonyl-CoA, which is further processed into palmitoyl-CoA and eventually fatty acids. (c) Schematic of selected genes with altered alternative splicing in DM1 livers, displaying locations of alternatively regulated exons and notable domains. Gene names are on the left. P = phosphorylation cluster; BC = biotin carboxylase domain; BB = biotin binding/biotin carboxylase carrier protein domain; CT = carboxyltransferase domain. (d) Percent inclusion of Acc1 exon 28 in chronic DM1 and controls and mice on a westernized diet (n=10, 13, 24, 21 from left to right). (e) Western blot showing Acc1 protein levels in chronic DM1 and control mice. (f) Lipid accumulation in acute and chronic DM1 mice livers via ORO staining. Black scale bars represent 50 µm. (g) Diagram summarizing gene abundance changes related to lipid regulation in livers of acute and chronic DM1 mice and chronic DM1 mice treated with Acc1 inhibitor CP-640816. Chronic DM1 mice transcripts compared to ApoE-rtTA mice fed 0.1g Dox for two months by RNA sequencing analysis. Acute DM1 and transcripts from DM1 mice treated with Acc1 inhibitor compared to chronic DM1 animals via qPCR. (h) RT-PCR splicing gel analysis of Acc1 exon 28 in acute and chronic DM1 mice livers. (i) Changes in Acc1 exon 28 inclusion in chronic DM1 mice treated with repeated doses of 12mg per kg body weight of either control antisense oligonucleotide (ASO) or Acc1 exon 28 targeting ASO. (j) ORO staining of chronic DM1 mice livers treated with Acc1 exon 28 or control ASO. Black scale bars represent 200 µm. (k) ORO staining from chronic DM1 mice treated with daily doses of 25µg of Acc1 inhibitor CP-640816 per kg of body weight or equivalent volume of vehicle. Black scale bars represent 200 µm. Values are displayed as median to quartiles for box plots. ***P < 0.001, ****P < 0.0001

The exon 28 that changes in the context of DM1 is centrally situated in the protein-coding region of the *Acc1* transcript; exclusion of the exon results in the loss of 24 amino acids directly N-terminal to the central phosphorylation cluster (Fig. 6c)^55,57^. This exon is included in *Acc1* transcripts in most tissues, having only been demonstrated to be excluded in the brain, mammary tissues, ovaries, and liver^57^. In mammary tissues of sheep and goats, skipping of this exon results in increased phosphorylation and decreased function of ACC1, altering the lipid profiles of ovine milk^57,58^.

In the livers of DM1 mice, exon 28 was included in ∼30% more *Acc1* transcripts than in control mice (Fig. 6a, d). Unexpectedly, MBNL1 deficiency in the liver did not affect *Acc1* splicing (Fig. 6a). Furthermore, upon feeding a high-fat, high-sugar diet, the inclusion of exon 28 increased slightly in control but not in DM1 mice, which maintained similar PSI values as on standard chow (Fig. 6d). Additionally, western blot analysis showed significantly higher ACC1 protein levels in the livers of DM1 mice relative to controls (Fig. 6e). Whether increased ACC1 protein abundance in DM1 mice livers is linked to exon 28 inclusion or occurred independently through another event regulating translation or protein stability remains to be determined.

To discern whether the upregulation of alternatively spliced ACC1 isoform was a primary consequence of DM1 or a secondary response to steatosis, we used an acute DM1 liver model. In contrast to the chronic model, “acute DM1” mice were aged for eight weeks before being fed 0.1g/kg Dox-containing diet for 12 days (Supplementary Fig. 3a). The acute DM1 mice did not exhibit any steatosis, which was evident in the chronic DM1 model (Fig. 6f). Next, we compared the hepatic mRNA abundance of genes that are directly linked to lipid biosynthesis, transport or metabolism-related functions in acute versus chronic DM1 mice via qPCR. In many cases, the mRNA levels were similar between acute and chronic DM1 livers. However, some transcripts in the acute model showed significant differences in the chronic model, highlighting transcripts that may have changed in response to pathological changes in the chronic DM1 liver (Fig. 6g, Supplementary Fig. 3b, c). Importantly, missplicing of *Acc1*, *Pnpla6*, and *Sorbs1* transcripts was evident in both acute and chronic DM1 liver models (Fig. 6h; Supplementary Fig. 3d). Acute DM1 livers also demonstrated significant increase in ACC1 protein levels; but, this increase was lower than in chronic DM1 livers (Supplementary Fig. 3b). These data demonstrate that ACC1 misregulation in DM1 is a direct effect of repeat RNA toxicity and not a secondary response to lipid accumulation or liver injury.

To determine if the upregulation of alternatively spliced ACC1 isoform contributes to DM1-related NAFLD, chronic DM1 mice were treated with antisense oligonucleotides (ASO) targeting the 5’ss of exon 28 of *Acc1* (Supplementary Fig. 3f). ASO treatment consisted of two loading doses of 12 mg/kg ASO the first two days of treatment, followed by two maintenance doses on days six and ten, and the mice were sacrificed on day eleven. After ASO treatment, the inclusion of exon 28 in DM1 mice livers dropped to less than 20%, an almost 60% decrease relative to untreated or control ASO-treated DM1 livers (Fig. 6i). Despite having a striking effect on *Acc1* splicing, the ASO treatment did not cause a noticeable change in ACC1 protein abundance (Supplementary Fig. 3e). These data indicate that increase in ACC1 and its splice isoform switch are likely two separate events and that upregulation of ACC1 protein in DM1 is not a consequence of higher protein stability or increased mRNA translation of the alternative isoform. We next evaluated the functionality of *Acc1* splicing redirection on lipid accumulation in DM1 mice livers. Chronic DM1 mice treated with ASO targeting *Acc1* exon 28 showed only a modest improvement in hepatic steatosis compared to control ASO treatment (Fig. 6j).

To further investigate whether ACC1 is required for the development of NAFLD phenotype in DM1-afflicted mice livers, we used the ACC1 inhibitor (CP-641086) to prevent ACC1 activity in these mice (Supplementary Fig. 4b). This inhibitor has been previously characterized for its ability to inhibit ACC1 activity, and it effectively reduces malonyl-CoA concentrations and fatty acid synthesis in cultured cells and mice livers^59,60^. We orally administered 25 μg/g ACC1 inhibitor twice daily to chronic DM1 mice for five days and then analyzed their liver health. Strikingly, short-term inhibition of ACC1 activity led to a significant recovery in hepatic steatosis in DM1-afflicted livers (Fig. 6k, Supplementary Fig. 4c) without affecting the expression of most lipid metabolism-related genes (Fig. 6j; Supplementary Fig. 3c). However, while the inhibition of ACC1 significantly reduced the appearance of fatty liver, markers of poor liver health associated with DM1-related NAFLD, such as increased hepatosomatic index and lobular inflammation failed to improve after treatment (Supplementary Fig. 4d, e). This suggests that these phenotypes in chronic DM1 mice either occur independently of ACC1 dysregulation or that a longer-term ACC1 inhibition is required to reverse the pathologies developing from sustained lipid accumulation. Collectively, these data demonstrate that upregulation of the alternative ACC1 splice isoform drives the bulk of hepatic lipid accumulation in DM1 and that short-term inhibition of ACC1 can reverse the hepatic steatosis phenotype in DM1-afflicted mice livers.

## Discussion

DM1 symptoms often extend beyond the musculature, wherein various tissues manifest specific pathologies that contribute to the affected individual’s overall health^10,26,61^. For instance, patients commonly experience gastrointestinal disturbances, metabolic dysregulation, increased sensitivity to drug injury, as well as the development of NAFLD and other liver dysfunctions^18–20,22,62–65^. However, the root cause of such dysfunctions is not fully understood. Elevated liver enzyme levels (GGT, AP) in DM1 individuals are also observed consistently but are speculated to occur secondary to the gallbladder and bile duct dysmotility, which might impair bile excretion and indirectly affect liver function^22,62^. In this study, we provide multiple lines of evidence that targeted expression of CUG repeat-containing RNA within hepatocytes is sufficient to alter the function of the liver, resulting in steatosis and hepatocellular injury. When combined with diet-induced metabolic stress, this predisposes the DM1 liver toward NAFLD and compromises its ability to respond to and metabolize specific analgesics and muscle relaxants. We determined that both NAFLD and poor drug metabolism phenotypes are direct consequences of repeat RNA toxicity in the liver, as these defects were not seen in the HSA L/R transgenic mice, which express repeat RNA only in the muscle tissues. Conversely, the HSA L/R mice suffer from significant glucose intolerance, whereas the DM1 liver mice are normal in glucose handling. These findings highlight the importance of studying the effects of DM1 within individual tissues and evaluating their respective contributions to the metabolic symptoms of this complex disease.

We further demonstrate that hepatic expression of toxic CUG960i RNA triggers global mRNA abundance and processing defects in genes enriched in ontologies that group into major functional clusters, especially lipid and drug metabolism, cell signaling and immune responses, as well as macromolecular binding and transport-related activities. These gene expression defects can directly lead to increased lipid accumulation and heighten the liver’s vulnerability to damage from toxins/dietary stress, partly explaining the high incidence of NAFLD and metabolic disorders seen in DM1 patients^19,20,22^. For instance, scores of genes involved in phase I and II drug metabolism were misregulated in the DM1 liver model. Notably, the levels of CYP2E1 and SULT2A1 proteins were significantly reduced in the livers of DM1 mice. A decrease in these two proteins alone may help to explain some of the poor responses to sedatives seen in DM1 patients, especially in the case of commonly inhaled anesthetics, which tend to be halogenated-hydrocarbons^66–68^. Further work is needed to determine whether abnormal drug metabolism or dysregulation of drug-metabolizing enzymes is a direct consequence of repeat RNA toxicity or an aberrant response to lipid accumulation and DM1-related NAFLD.

The pathological mechanism of DM1 involves sequestration and disruption of MBNL protein activities, shifting the affected tissues’ transcriptome from an adult-to-preadolescent-like state. Also, MBNL expression is upregulated in hepatocytes as the liver matures after birth^69,70^. Intriguingly, when we compared the hepatic transcriptomes of DM1 and *Mbnl1* KO mice, less than a third of the misregulated events occurring in the DM1 hepatocytes were detected in MBNL1 deficient hepatocytes. The dissimilarities between the DM1 liver model and *Mbnl1* KOs were also evident at the tissue level, as many of the pathological consequences of expressing DM1 in the liver were not seen in the *Mbnl1* KO mice, including the lack of lipid accumulation. MBNL proteins serve critical functions in maintaining tissue maturity; however, which MBNL family member is predominantly responsible for these activities is tissue-dependent^11,71–74^. Additionally, MBNL proteins have high sequence and structure similarity and significant overlap in binding targets; thus, substitution between the two predominant members, MBNL1 and MBNL2, allows for a tunable regulatory system that maintains appropriate splicing form for the bulk of co-regulated transcripts^73–75^. Our examination of *Mbnl1* KO mice revealed that MBNL2 protein levels are increased in the KO livers, a phenomenon also seen in other tissues^34,72,74,76^. In the future, it would be interesting to determine whether elevated expression of MBNL2 could explain the lack of overlap in transcriptome changes between the *Mbnl1* KO and DM1 livers, as well as shed light on possible compensatory mechanism(s) that buffer the effects of MBNL1 loss within the liver.

Several genes associated with insulin-regulated lipid metabolism were altered in DM1-afflicted livers. Because lipid homeostasis is regulated jointly by intra- and extra-hepatic signaling, these changes could be a primary result of repeat RNA toxicity or a secondary effect of the diseased liver. Indeed, there is some evidence of a secondary effect occurring — the expression and/or splicing of genes such as *Sorbs1*, *Fabp1*, *Acox1*, and *Hsd17b13* were altered less in the acute DM1 liver as compared to the chronic DM1 liver, possibly indicating effects that are exacerbated with lipid accumulation. However, based on our analysis, missplicing and upregulation of ACC1 is a rapid response to the repeat RNA expression. ACC1 is a focal point for *de novo* lipid biosynthesis, converting excess acetyl-CoA into malonyl-CoA, which can then be used to produce palmitate^77,78^. Therefore, ACC1 activity must be tightly controlled in response to the nutritional state of the liver as misregulation of ACC1 stimulates excess lipogenesis and NAFLD phenotypes^79,80^.

Our results indicate that short-term inhibition of ACC1 activity is sufficient to reverse lipid accumulation in the DM1-afflicted mice livers, highlighting the importance and involvement of this pathway. We believe this beneficial effect is primarily due to inhibition of *de novo* lipogenesis; however, reduced production of malonyl-CoA would also increase the mitochondrial import and oxidation of fatty acids. Although extrapolating these findings to DM1-related NAFLD in humans is not straightforward, pharmacologic inhibition of ACC is currently viewed as one of the most promising therapeutic approaches for treating NAFLD/NASH^81–85^. It is important to note that a small proportion of NAFLD patients treated with ACC inhibitors may experience hypertriglyceridemia over time, which can increase their risk of developing cardiovascular disease^86–88^. Whether inhibiting ACC1 in the context of DM1 leads to increased serum triglycerides remains to be determined. Therefore, in future studies, it will be essential to examine the long-term effects of ACC1 inhibitors on systemic lipid trafficking and mobilization in DM1 liver mice.

In conclusion, our study provides the first characterization of the direct impact of DM1 on liver health. The findings offer valuable insights into how disrupted hepatic functions contribute to the metabolic symptoms and drug sensitivities in DM1, underscoring the idea that a malfunctioning liver can further complicate the treatment of this complex genetic disease. A current limitation of this work is that our results need to be validated in humans, and the extent of similarity between the murine DM1 liver model and actual patient livers must be systematically assessed. Therefore, enhanced screening for NAFLD/hepatocellular injury in affected individuals, along with access to liver biopsies, is urgently needed because if DM1 livers cannot provide adequate metabolism of xenobiotic material, it would prolong the clearance time for many drugs, altering their therapeutic index. Future investigations incorporating patient samples and combining clinical data with the mouse model findings will be pivotal in determining the extent of hepatic dysfunctions in DM1. This will help ensure the applicability, optimization, and effectiveness of prospective treatments being developed to treat/manage various symptoms of this debilitating disease.

## Methods

### Development of the DM1 Liver Model

The “DM1 liver” line was generated by crossbreeding TRE-960i mice with ApoE-rtTA mice. ApoE-rtTA mice express a reverse tetracycline transactivator (rtTA) under a liver-specific ApoE promoter^89^. TRE-960i mice carry a tetracycline response element (TRE)-driven truncated DMPK gene with a (CUG)_960_ repeat sequence in the final exon^23^. Mice with ApoE-rtTA alone were used as controls. Both lines were maintained as homozygotes for transgenic alleles.

Mouse care and use followed NIH guidelines for animal care, and animal protocols were approved by the Institutional Animal Care and Use Committee at the University of Illinois at Urbana-Champaign.

### Mouse diet schemes

In most experiments, DM1 liver and ApoE-rtTA mice were subjected to the disease-inducing Dox diet from birth, with mothers receiving 2.0 g/kg Dox-supplemented Teklad 2018 18% protein global rodent diet until weaning at 21 days. Subsequently, weaned mice transitioned to a 0.1 g/kg Dox diet until sacrifice at nine weeks. This feeding protocol is referred to as the chronic DM1 liver model.

Exceptions to the protocol above include: DM1 liver mice used for RNA-seq, which remained on a 2 g/kg Dox diet until sacrifice. “No-Dox” mice were ApoE-rtTA; TRE960i mice maintained on a Dox-free diet. “Recovery” mice followed the chronic DM1 protocol but were transitioned to a Dox-free diet for ten days before sacrifice. For the Western Diet model, mothers were on 2g/kg Dox-supplemented chow until weaning. Then, mice were switched to a high-fat, high-sugar, cholesterol-supplemented “Western” diet (Teklad 88137) supplemented with 0.1g/kg Dox for eight weeks. In the acute DM1 model, mice were Dox-free until eight weeks, followed by a 0.1 g/kg Dox diet for 12 or 18 days before sacrifice.

### Zoxazolamine recovery testing

Zoxazolamine (Zox) solution, prepared the day before, consisted of Zox dissolved in DMSO to achieve a final concentration of 15 µg/µL in a 95% Corn-Oil, 5% DMSO solution. After an 18-22 hour fast, mice received 120 mg/kg Zox injections. Vigorous homogenization between injections ensured a uniform solution. Post-treatment, mice freely roamed until they lost motor function, after which they were placed supine on an insulating blanket. Time was recorded until mice successfully self-righted three times^90^.

### APAP insult testing

Acetaminophen (APAP) solutions were freshly prepared, dissolving 20 mg of APAP in 1 mL sterile 1x PBS. After heating at 55°C for 15 minutes with periodic vortexing, the solution was maintained at 40°C during injection, with thorough mixing between injections. Mice fasted for 18-22 hours and received a 350 mg/kg IP injection of APAP. Mice were observed for 8 hours when surviving mice returned to the animal care facility. Harvesting of serum and liver samples occurred 24 hours post-APAP injection^91,92^.

### *Acc*1 exon 28 ASO treatment

*Acc1* Exon 28 ASO, developed by Gene Tools, LLC, targeted the 5’ splicing site of *Acc1* exon 28 (sequence: CCCTCTGTAATTAAA). A standard control in-vivo morpholino with the sequence CCTCTTACCTCAGTT served as a control. ASOs were dissolved in sterile biology-grade water. Mice designated for ASO treatment followed the chronic DM1 model until day 63, receiving IP injections of 12 mg ASO per kg body weight on days 63, 64, 68, and 72. Mice were sacrificed after a 4-6 hour fast on day 73.

### ACC1 inhibitor treatment

ACC1 inhibitor CP-641086, procured from MedChemExpress (Lot#12439), was dissolved in a 5% (w/w) methylcellulose solution in molecular biology-grade water. Mice selected for ACC1 inhibitor treatment followed the chronic DM1 model until day 63. They received doses of either 25 µg inhibitor or vehicle (5% methylcellulose) per g body weight twice daily via oral gavage for five days. Sacrifice occurred on the sixth day after 4-6 hours of fasting.

### Statistical analysis and data visualization

All quantitative experiments have at least three independent biological repeats. The results were expressed with mean and standard deviation unless mentioned otherwise.

Differences between groups were examined for statistical significance using unpaired T-tests when comparing directly between groups or one-way analysis of variance (ANOVA) for more than two groups using the GraphPad Prism 9 Software. Statistical outliers were determined with the ROUT method in Prism, with Q = 5%. P-value < 0.05 or FDR < 0.10 was considered significant. RNA-seq data plots were generated in R using the ggplot2 package. In all figures, significance was set as p < 0.05, “*” indicates p < 0.05, “**” indicates p < 0.01, “***” indicates p < 0.001, and “****” indicates p < 0.0001. Data presented as bar graph or linear For Box plots: median is represented as center line, median; box limits are the upper and lower quartiles; whiskers are set as 1.5x interquartile range.

## RESOURCE AVAILABILITY

### Data Availability

RNA-Seq data that support the findings of this study have been deposited in NCBI Gene Expression Omnibus under the primary accession code GSE252827 (https://www.ncbi.nlm.nih.gov/geo/query/acc.cgi?acc=GSE252827)

### Materials Availability

Requests for reagents, resources, and additional information should be directed to the corresponding author, Auinash Kalsotra.

## Acknowledgments

We acknowledge support from the Transgenic mouse core, High-throughput sequencing and genotyping core, and Histology-microscopy core facilities at the University of Illinois, Urbana-Champaign. We thank the members of the Kalsotra and Anakk laboratories for their valuable discussions and comments on the manuscript.

## Author Contributions Statement

Z.D. and A.K. conceived the project and designed the experiments. Z.D. and A.G. performed experiments and analyzed the data. Z.D., AK, and U.V.C facilitated mouse model development and management. Z.D. and A.K. interpreted the results and wrote the manuscript. All authors discussed the results and edited the manuscript.

