## Supplemental Files for "Altered drug metabolism and increased susceptibility to fatty liver disease in myotonic dystrophy"

#### Supplementary Figures

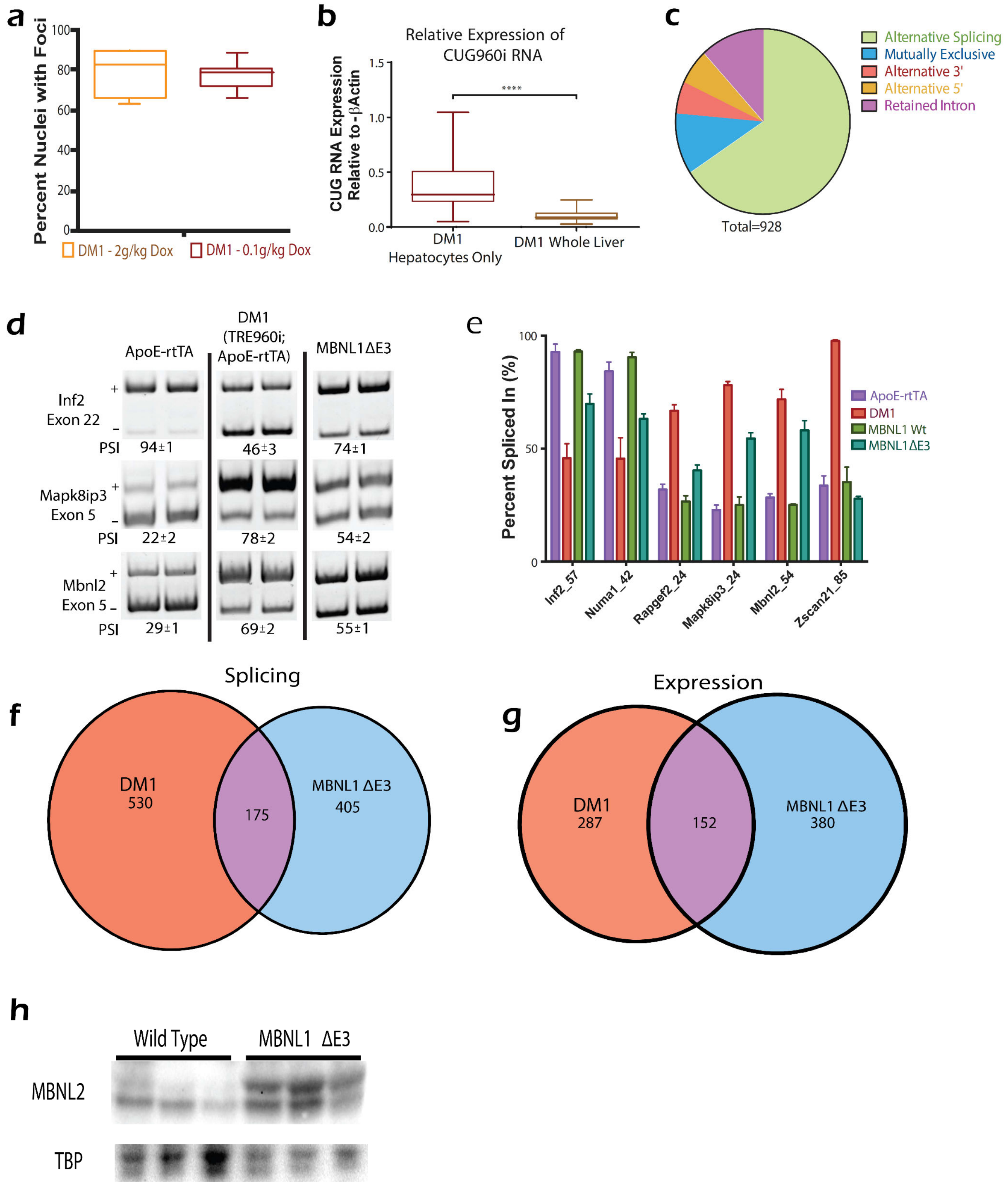

**Supplemental Figure 1: Comparison of Hepatic Transcriptome Changes in DM1 Models.**

(a) RNA FISH-IF quantification of CUG960i/Mbnl1 foci in hepatocyte nuclei for mice on 2g/kg Dox diet vs. 0.1 g/kg Dox diet (n=7 per group). (b) Quantitative-PCR analysis of toxic CUG960i

RNA in DM1 liver mice hepatocytes (n=3) and whole livers (n=20). (c) Pie chart representing categories of alternative splicing events from RNA-seq data. (d) RT-PCR splice assay for selected MBNL1 targets, indicating exon inclusion (+) or exclusion (-) with PSI listed below. (e) PSI quantification for events changing in MBNL1 KO or DM1 liver mice, compared to controls (n≥6 for ApoE-rtTA mice, n≥6 for DM1 liver mice, n≥3 for MBNL1 WT mice and n≥4 for MBNL1 KO mice) (f) Venn diagram comparing alternative splicing events in DM1 liver or MBNL1 KO models relative to controls. (g) Venn diagram comparing DEG in DM1 liver or MBNL1 KO models relative to controls. (h) Western blot showing increased MBNL2 protein in MBNL1 KO mice. Values displayed as median to quartiles for box plots; mean ± SD for all others. ****P < 0.0001.

| Cluster Name | Example Ontologies Effected | Total Genes Affected | Enrichment Score | P-Value |
|---|---|---|---|---|
| **ZINC FINGER PROTEINS** | IPR001965: Zinc Finger, PHD-Type | 12 | 3.375 | 6.14E-5 |
| **KINASE AND PHOSPHORYLATION ACTIVITY** | | | 3.166 | |
| | GO:0000166: Nucleotide Binding | 92 | | 3.63E-5 |
| | GO:0006468: Protein Phosphorylation | 36 | | 7.77E-5 |
| | IPR008271: Serine/threonine-Protein Kinase | 23 | | 2.89E-4 |

**Supplemental Table 1: Gene Ontology Clusters: Alternative Splicing**. Gene ontology clusters identified by DAVID Functional Annotation within transcripts undergoing alternative splicing in DM1-afflicted hepatocytes.

| Cluster Name | Example Ontologies Effected | Total Genes Affected | Enrichment Score | P-Value |
|---|---|---|---|---|
| **Cytochrome p450** | | | 6.286 | |
| | GO:0020037: Heme Binding | 31 | | 3.95E-14 |
| | IPR002401: Cytochrome P450, E-class, Group 1 | 18 | | 3.29E-10 |
| | GO:008392: Arachidonic Acid Metabolism | 18 | | 1.55E-9 |
| **Kinase and Phosphorylation activity** | | | 2.97 | |
| | GO:0006468: Protein Phosphorylation | 40 | | 7.30E-6 |
| | IPR001245: Serine/threonine-Protein Kinase | 13 | | 9.87E-4 |
| **SMAD Protein Pathways** | GO:060389-SMAD protein phosphorylation | 5 | 2.89 | 5.64E-4 |
| **Hormone Receptor Signaling** | | | 2.89 | |
| | GO:0003707: Steroid Hormone Receptor Activity | 9 | | 3.69E-4 |
| | IPR000536: Nuclear hormone receptor | 8 | | 6.03E-4 |
| **C-type Lectin Domains** | IPR016186:C-type lectin-like | 13 | 2.48 | 8.69E-4 |

**Supplemental Table 2: Gene Ontology Clusters: Increasing Abundance**. Gene ontology clusters identified by DAVID Functional Annotation within DEGs increasing in abundance in DM1-afflicted hepatocytes.

| Cluster Name | Example Ontologies Effected | Total Genes Affected | Enrichment Score | P-Value |
|---|---|---|---|---|
| **Hormone Biosynthesis** | mmu00830: Retinol metabolism | 7 | 2.48 | 1.69E-3 |
| **Regulation of Development** | | | 2.41 | |
| | mmu04950:Maturity onset diabetes of the young | 6 | | 3.91E-5 |
| | GO:0001889~liver development | 5 | | 1.49E-3 |

**Supplemental Table 3: Gene Ontology Clusters: Decreasing Abundance**. Gene ontology clusters identified by DAVID Functional Annotation within DEGs decreasing in abundance in DM1-afflicted hepatocytes.

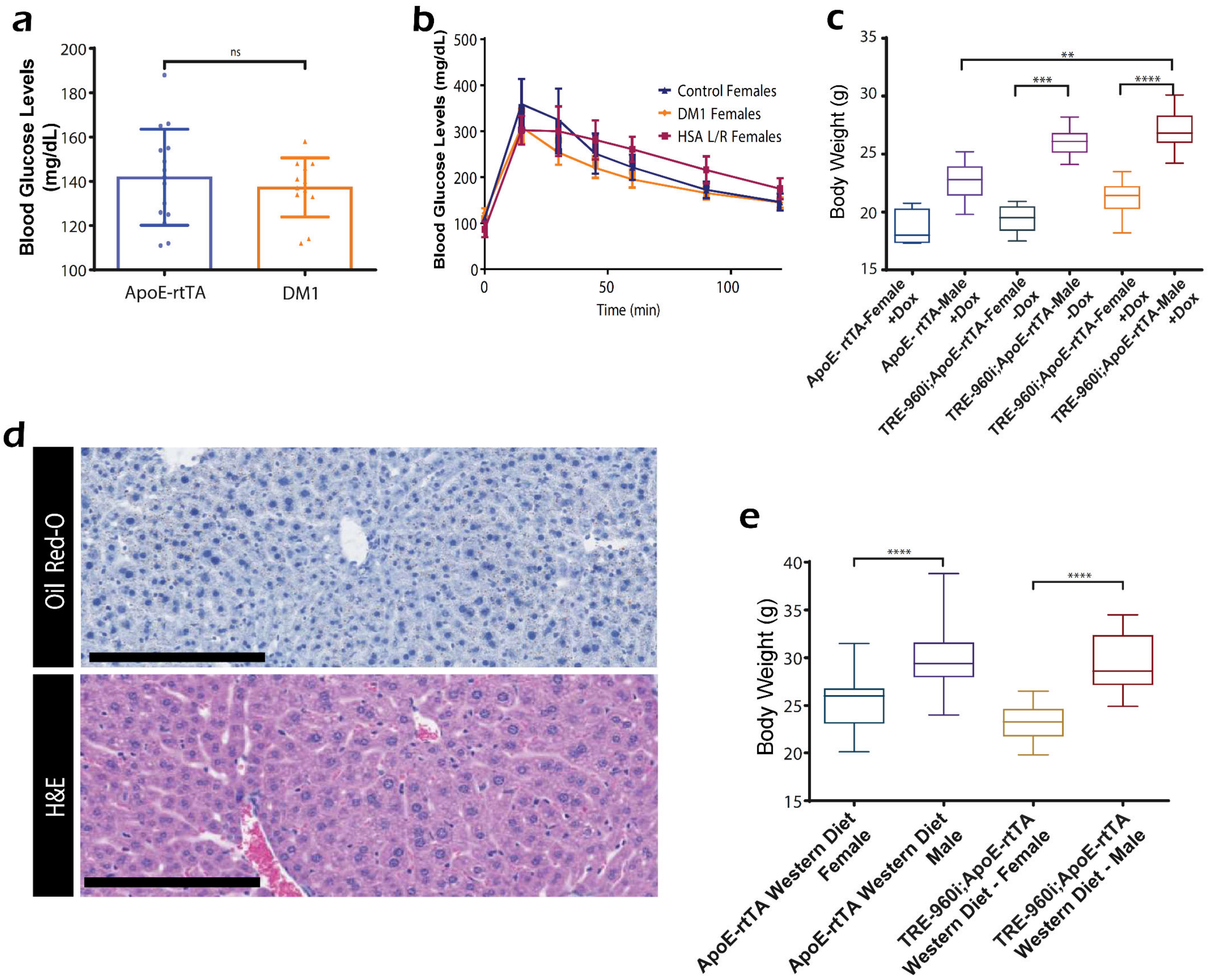

**Supplemental Figure 2: Physiological Changes Across Sex, Diet, and Models.** (a) Blood glucose levels in fasting female mice (n=15 for ApoE-rtTA and 12 for DM1 liver mice). (b) Glucose tolerance testing (GTT) curves in female mice after IP glucose injection (n=5 ApoE-rtTA, 14 DM1 liver, 9 HSA L/R mice). (c) Mean body masses of mice at sacrifice (n=9 male ApoE-rtTA, 5 female ApoE-rtTA, 9 DM1 male w/o Dox, 5 DM1 female w/o Dox, 10 DM1 liver male, 9 DM1 liver female). (d) Representative images of HSA L/R mouse livers - ORO (top) and H&E staining (bottom). (e) Mean body masses of mice on a western diet at sacrifice (n=17 male and female ApoE-rtTA, 10 DM1 liver male, 9 DM1 liver female mice). Values displayed as median to quartiles for box plots; mean ± SD for all others. $^{**}P < 0.01$, $^{***}P < 0.001$, $^{****}P < 0.0001$.

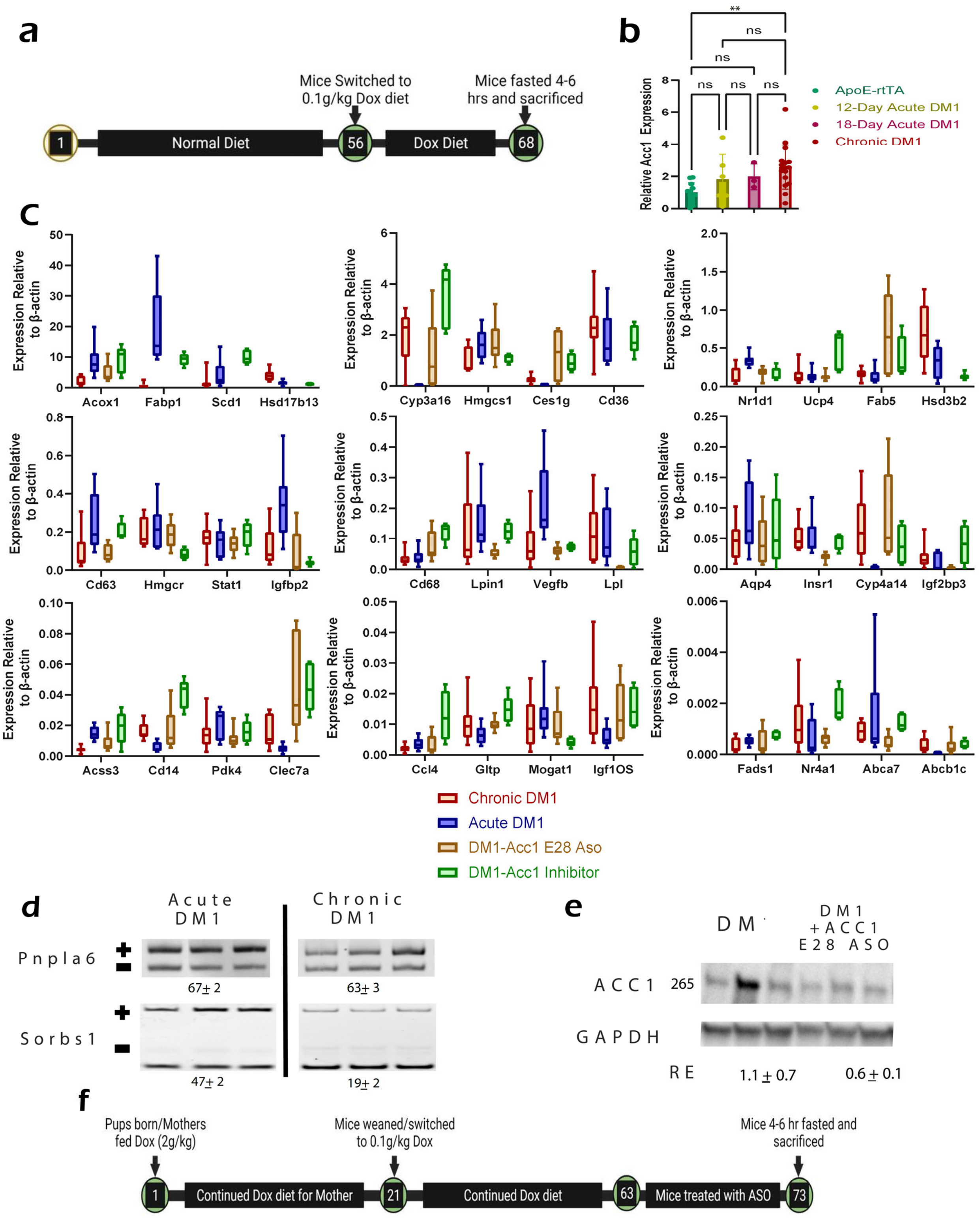

**Supplemental Figure 3: Ancillary Data for Acute DM1 Models and Mice Treated with *Acc1* ASO** (a) Diet program schematic for acute DM1 mice: standard chow until eight weeks, then switched to 0.1g/kg. (b) ACC1 protein levels, as determined by western blot, in control, 12-day and 18-day acute DM1, and chronic DM1 models (n=12, 6, 3, and 16, left to right). (c) Bar graphs of qPCR analyses for Fig. 6H ($n \geq 5$ for acute DM1, $n \geq 8$ for chronic DM1, $n \geq 7$ for *Acc1* E28 Aso treated, and $n \geq 4$ for ACC1 inhibitor treated). (d) RT-PCR splicing gel analysis of Pnpla6 exon two and Sorbs1 exon twenty-three in acute and chronic DM1 mouse livers. (e) Western blot analysis of Acc1 in chronic DM1 mice vs. those treated with anti-*Acc1* exon 28 ASO. (f) Diet and treatment schedule schematic for chronic DM1 mice treated with *Acc1* exon 28 ASO or control ASO. Box plots show the first to third quartile with a median line; values as mean ± SD for all others. **P < 0.01.

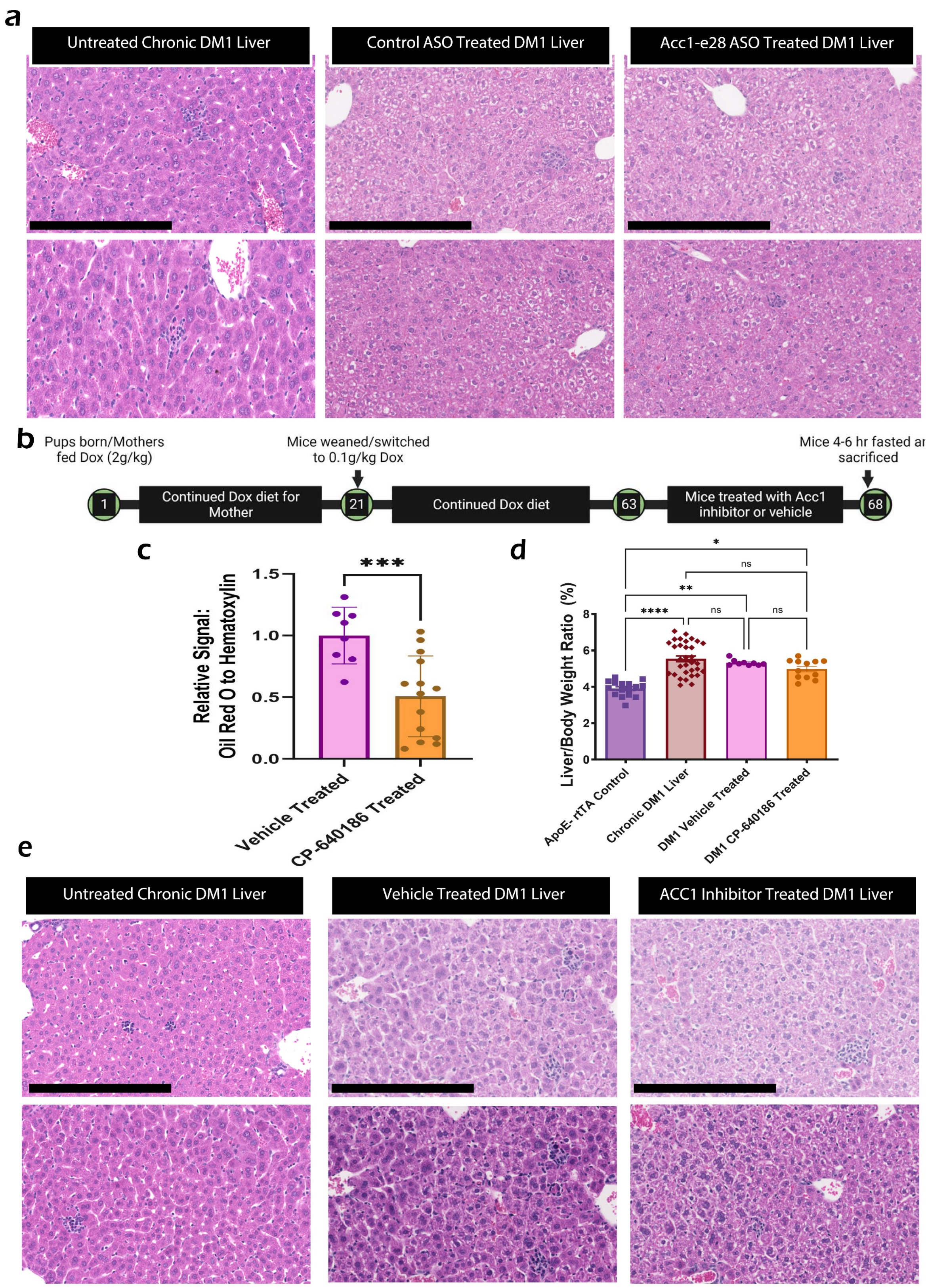

**Supplemental Figure 4: Physiological analysis of DM1 mice treated with *Acc1 e28* ASO and ACC1 Inhibitor Treated Animals**. (a) Representative H&E images of chronic DM1 mice and mice treated with control or anti-*Acc1* exon 28 ASO. Black scale bars represent 250 µm. (b) Feeding and treatment protocol schematic to inhibit ACC1 function in chronic DM1 mice. Mice receive ACC1 inhibitor CP-640816 twice daily from nine weeks of age, sacrificed on the sixth day after a four-hour fast. (c) Quantification of Oil Red O signal, relative to hematoxylin-stained nuclei, indicating lipid accumulation chronic DM1 liver mice treated with ACC1 inhibitor CP-640816 or vehicle (n=16, 32, 9, and 12, left to right). (d) Hepatosomatic index for chronic DM1 mice and respective controls compared to chronic DM1 mice treated with ACC1 inhibitor CP-640816 or vehicle (n n=8 for vehicle-treated and 14 for CP-640816 treated mice) (e) Representative H&E images of livers from chronic DM1 mice treated with ACC1 inhibitor CP-640816 and vehicle. Black scale bars represent 250 µm. Values represent mean ± SD. *P < 0.05, **P < 0.01, ***P < 0.001, ****P < 0.0001.

| Antibody | Host Species | Type | Application | Supplier Information |
|---|---|---|---|---|
| MBNL1 | Mouse | Primary | IF – 1:500<br>WB – 1:1000 | Santa Cruz Biotechnology, sc-47740 |
| MBNL2 | Mouse | Primary | IF – 1:500<br>WB – 1:1000 | Santa Cruz Biotechnology, sc-136167 |
| anti-Mouse, Dylight 488 | Goat | Secondary, Fluorescence | IF – 1:500 | Thermo Fisher Scientific, AB_1965946 |
| anti-Mouse, Dylight 594 | Goat | Secondary, Fluorescence | IF – 1:500 | Thermo Fisher Scientific, AB_1965950 |
| TBP | Mouse | Primary | WB – 1:5000 | Thermo Fisher Scientific, AB_10980989 |
| anti-Mouse | Goat | Secondary, HRP | WB – 1:5000 | Bio-Rad, #1706516 |
| CYP2E1 | Rabbit | Primary | WB – 1:500 | Abcam, ab28146 |
| SULT2a1 | Rabbit | Primary | WB – 1:500 | Abcam, ab194113 |
| GAPDH | Rabbit | Primary | WB – 1:5000 | Cell Signaling Technology, #2118 |
| anti-Rabbit | Goat | Secondary, HRP | WB – 1:5000 | Thermo Fisher Scientific, AB_228341 |
| ACC1 | Rabbit | Primary | WB – 1:500 | Cell Signaling Technology, #4190 |

**Supplementary Table 4. Antibody dilutions and supplier information.** WB = western blot, IHC = immunohistochemistry, IF = immunofluorescence.

| Gene | Forwards Primer | Reverse Primer |
|---|---|---|
| β-Actin | CCCTAAGGCCAACGGTGAAA | CGGAGTCCATCACAATGCCT |
| TRE-*960i* | GGGCCGTCCGTGTTC | GGGCGTCATGCACAAGAAAG |
| Abca7 | CACAGAGAAGGTGGGGACTC | AAGCCCTGTGGGGCCAG |
| Abcb1a | CGGAGTCAGACAGAACAAGAAGA | CTTCTTACTCCATTCCCCCTTT |
| Abcd2 | GCGGATGTTTTACCATAAACCG | TAGGGAAATCCCAGCCCCAA |
| Acox1 | AGCTACGTGCAGCCAGATTG | AGAAGTCAAGTTCCACGCCA |
| Acss3 | AGAGACTGGATCCCCCATCA | TGAAGCAGGTATCGTCCATTGTA |
| Aqp4 | CTGGGCAAACCACTGGATATATTG | TGAGCTCCACATCAGGACAG |
| Avpr1a | CTACATCCTCTGCTGGACACC | GGAAGGGTTTTCGGAATCGGT |
| Ccl4 | AACCTAACCCCGAGCAACAC | AGGGTCAGAGCCCATTGGTG |
| Cd14 | AGAATCTACCGACCATGGAGC | ACTTTCCTCGTCTAGCTCGC |
| Cd36 | TTAATGGCACAGACGCAGCC | TCAGATCCGAACACAGCGTAG |

| | | |
|---|---|---|
| Cd63 | CTATCCATACCCAGGGCTGC | CCTCCACAAAAGCAATGCCC |
| Ces1g | ACCTCACTAGACCCCAGAGC | AGGATGGGTGTCCCAGACTC |
| Cyp3a16 | TTTTGTGGAGAATGCCAAGAAGG | TGAGGAATGGAAAGAGTGCTAC |
| Cyp4a10/31/32 | TGGGGCGATCAGATCCAAAG | TGGGGTTAGCATCCTCCTGT |
| Cyp4a14 | GGCACCATCTGAAGGACAAG | GCTCCCCGAGAGACACTGTA |
| Elovl2 | ATACCTTGTGGTCAAAGCTTCTT | GTACTTGTGCATGGACGGGA |
| Elovl3 | CTACACGGATGACGCCGTAG | ATGAAGGCCGTGTCTCCCAG |
| Fabp1 | CTACACGGATGACGCCGTAG | ATGAAGGCCGTGTCTCCCAG |
| Fabp5 | TCCCACCATGGCCAGTCTTA | TAAGAGCCAGTCCTACTCCTAGC |
| Gltp | GCCGCCCTTCTTTGATTGC | GGTCGGTGTCATATACGGCT |
| Hmgcr | ATCCTGACGATAACGCGGTG | AAGAGGCCAGCAATACCCAG |
| Hmgcs1 | GCCCCTTCACAAATGACCAC | GCAGGGCTTGGAATATGCTCT |
| Hsd17b13 | ACAACTCTGTGGATCAGGTAAAGA | TCTCCTCGTCCTTGGCACTA |
| Hsd3b2 | ATCAGGGTCCTGGACAAGGT | TGATGCTTGTCTCTAGGTTGAAGA |
| Igf1os | TTCCAGGAAGGCCCCTAAGA | TCACCGCTGAGTTCCTTGTG |
| Igf2bp3 | ACGCTAAGATCCCGGTGG | CCCATGTAGTTCCATTTTACCTGA |
| Igfbp2 | GCCCCCTGGAACATCTCTAC | TCAGAGACATCTTGCACTGCT |
| Lpin1 | CACAGGCTGCAAAGTCCTCT | TCGCTGTGAATGGCCTGAAA |
| Lpl | AGGACTCAGCAGTGTTTGTGA | TTTGTTTGTCCAGTGTCAGCC |
| Lrp11 | CCCACAGACGGGGTAGTTCT | CTGGCTGAGGCACCTTCAT |
| Ly6a | GAAACCCCTCCCTCTTCAGG | GCTGCACAGATAAAACTTCCTCTC |
| Ly6d | AAAACCGTCACCTCAGTGGAG | CAGCATTGTGTGACCTCGGA |
| Mogat1 | CTGGAGAACCCCAGAGCAAG | CCTGCGTTTTGACAAGACAGAT |
| Nr1d1 | ATGTCTAGAGATGCTGTGCGT | GTCTCTAGAGGGCACAGGCT |
| Nr4a1 | GAAAGTTGGGGGAGTGTGCTA | TTGAATACAGGGCATCTCCAG |
| Pdk4 | AATGCCCCTTTGGCTGGTTT | AGCGTCTGTCCCATAACCTG |
| Scd1 | CCTACACCAACGGGCTCC | TGTAAGAACTGGAGATCTCTTGGA |
| Stat1 | CAGACCCACTTGGGACACTG | GAGCTGAAACGACTGGCTCT |
| Ubd | CTGTCCGCACCTGTGTTGT | GAGACCTTGGTTTGGGACCTA |
| Vegfb | CCAAGTCCGAATGCAGATCCT | TTGGCTGTGTTCTTCCAGGG |
| Vldlr | GTCCTGGAGATGCGATGGTG | CACTACATGTTATGTTGCCACAGTT |
| Wfdc2 | CTTTGGACAAGGACTGTGCG | GGTGCCCTGCTTTTCATTAGG |

**Supplementary Table 5.** Primer Sequences for qPCR analysis

| Gene | Forwards Primer | Reverse Primer |
|---|---|---|
| Git2 | GATCAGGCCAGAGCCATAC | AGGGTCACAGGCAGTGTTGT |
| Inf2 | GGATGAGGATTGAGAGGACA | GAGCACTCACTTGGCTTTGG |
| Mapk8ip3 | CGGCACACAGAGATGATCCA | TGTGGTACTGGGTGTTGCAG |
| Mbnl1 | GCTGCCCAATACCAGGTCAAC | TGGTGGGAGAAATGCTGTATGC |
| Mbnl2 | ATTTTCACCCTGCTGGACCAC | TTTGGTAAGGGATGAAGACCA |
| Myo1b | ACAAAGCGGTACCAGCAGA | TGCGTACCTTCAGTCCAAGC |
| Acc1 | TGGCAGCTCTGGAGGTGTATG | GACTCTGGGAATGTGGGGCTTT |
| Pnpla6 | TTTGCCCCGGATCGATTTGT | TGACCTAGCACCCGAACATT |
| Sorbs1 | TCAGAGTCACCAAGACATTTTATACC | ATTGGCTGGAGCAGGTCT |
| Nr1h4 | AGGGGATGAGCTGTGTGTTG | AGTTCCGTTTTCTCCCTGCAA |

**Supplementary Table 6.** Primer Sequences for RT-PCR Splice Assays

### Supplementary Methods

#### *Mouse handling and care*

National Institutes of Health (NIH) and University of Illinois, Urbana-Champaign (UIUC) institutional guidelines were followed in using and caring for laboratory animals. All experimental protocols were performed as approved by the Institutional Animal Care and Use Committee. The study is not gender-specific, and specimens include both male and female animals; however, as attributes such as glucose regulation and body weight are sex-specific, animal sex was recorded. Whole liver tissues and hepatocytes were isolated from mice following guidelines for euthanasia and anesthesia.

#### *Animal Models*

Four mouse models were utilized in this study. First, the control animals used for the DM1 experiment, the ApoE-rtTA mice, were a mixed strain C57Bl6/DBA mouse line, with a single transgene containing a reverse tetracycline TransActivator (rtTA), expressed under the ApoE promoter[1]. Second, the "DM1 liver" line was also a mixed strain line, combining FVB background TRE-960i mice with the ApoE-rtTA mice. The resulting FVB/C57Bl6/DBA mice contained both the ApoE-rtTA transgene as well as tetracycline response element (TRE) driven truncated DMPK gene containing only the last five exons of human DMPK [2]. The DMPK construct also contained an elongated CUG repeat sequence with 960 repeats. These repeats are interrupted every twenty repeats with a "ctcga" sequence to prevent the overall repeat sequence from undergoing expansion or shrinkage. The ApoE-rtTA and DM1 liver mice were maintained as homozygotes for all transgenic alleles.

HSA L/R mice were FVB mice expressing a truncated human skeletal actin (HSA) with a ~240 CUG repeat sequence in the 3' UTR of the transgene[3]. The HSA was driven by the skeletal actin promoter, allowing the exclusive expression of CUG repeats within the skeletal muscle tissue. The HSA L/R mice were maintained as homozygotes.

The final model, the *Mbnl1* knock out (KO) or *Mbnl1*$^{\Delta E3/\Delta E3}$ line, was an FVB mouse where the first coding exon of *Mbnl1* was replaced with a cassette using cre-lox insertion [4]. This mouse line was maintained in the heterozygous state, and homozygous mutant (*Mbnl1*$^{\Delta E3/\Delta E3}$) or homozygous wildtype (*Mbnl1*$^{Wt/Wt}$) were generated for study or as controls when needed.

*Diet schemes*

For most experiments performed in this study, DM1 liver and ApoE-rtTA mice were fed under the following scheme. To mimic the DM1 conditions in human patients, we induced the disease at birth, giving the mothers 2.0 g/kg doxycycline (Dox) supplemented Teklad 2018 18% protein global rodent diet. The 2.0 g/kg Dox diet continues until weening, 21 days after birth. The diet of the mice was then switched to a 0.1 g/kg Dox-supplemented Teklad 2018 global rodent diet and maintained on this diet until sacrificed at nine weeks of age.

Some important exceptions were the mice used for RNA-seq, which were maintained on a 2 g/kg Dox-supplemented diet until sacrifice at nine weeks. This model is referred to as the chronic DM1 model. Mice noted as “No-Dox” were fed only a global rodent diet without Dox until sacrifice at nine weeks. Mice noted as “Recovery” were fed 0.1g/kg Dox-supplemented diet as per the chronic DM1 model but then switched to a Dox-free diet for ten days before being sacrificed at nine weeks of age.

Western Diet mice were fed as before, with the mothers being fed 2g/kg Dox supplemented chow until weening, at which point the mice were transitioned to a high fat, high sugar, cholesterol supplemented “western, purified atherogenic” Diet (Teklad 88137), supplemented with 0.1g/kg Dox. These mice were maintained on this diet for eight weeks after weening until sacrificed at 11 weeks.

For the acute DM1 model, mice were fed a global rodent diet without Dox until eight weeks of age, after which mice were switched to a 0.1 g/kg Dox-supplemented diet. Mice were fed this diet for 12 or 18 days, after which mice were sacrificed.

*Glucose tolerance testing (GTT)*

GTT was performed on male and female mice, either seven days before sacrifice if they were maintained on the 0.1g/kg Dox diet or 10 and 5 days before sacrifice if they were on the Western diet. GTT was performed after the mice were fasted for 24 hours for both cases. Glucose was injected through intraperitoneal injection (IP) at a 2 g/kg body weight concentration. Tails were clipped, blood was collected, and glucose was measured using a One Touch Ultra 2 glucose meter after 0, 15, 30, 60, 90, and 120 minutes.

*Sacrifice and tissue collection*

Mice were fasted in the morning and were sacrificed after 4 to 6 hours of fasting. Liver and carcass weight were taken at the time of sacrifice, as well as 600 µL of blood via retro-orbital bleeding. During sacrificing, liver tissues were collected for (1) RNA, protein, and lipid isolation, (2) paraffin-embedding, and (3) cryo-sectioning. Tissue for cryo-sectioning was collected by sectioning two pieces of liver, embedding the tissues in Optimal Cutting Temperature (OCT) compound and frozen on dry ice. Tissues for paraffin embedding were stored in neutral buffered formalin for 48 hours before being stored in 70% ethanol until paraffin embedding. The remaining tissue was flash-frozen in liquid nitrogen.

*Isolation of hepatocytes*

Hepatocyte isolation was performed using two-step perfusion with centrifugal separation to produce a cell population highly enriched in hepatocytes; this population was then used for RNA-seq analysis. The method for hepatocyte isolation was adapted from Li et al., 2010.

Briefly, mice were anesthetized in a chamber supplied with isoflurane and oxygen (2.5% isoflurane in oxygen, 1.5Lmin.). Mice were maintained on the anesthetic during the procedure using a nose cone. The liver was perfused via cannulation of the portal vein with 30-40 ml of a 1× HBSS (Hank's balanced salt solution) with phenol red (without $Ca^{2+}$ and $Mg^{2+}$), 0.5 mM EDTA solution at a flow rate between 3 and 5 mL per minute. This solution was followed by 50 ml of a 1× HBSS (with Ca2+), 5.4 mM CaCl2, 0.04 mg ml−1 soybean trypsin inhibitor, and 3000 units of collagenase type I (Worthington Chemicals). Subsequently, the liver was massaged in a Petri dish containing 1× HBSS with phenol red (without $Ca^{2+}$ and $Mg^{2+}$) to release cells from the liver capsule, and then the cell suspension was passed through a 70-μm filter to obtain a single-cell suspension. The cells were centrifuged at 50 × g for 5 min (4 °C) to separate live hepatocytes from non-parenchymal and dead cells. The cells were washed three times in 1× HBSS as above, then flash-frozen in liquid nitrogen and stored at −80 °C until further use.

*Total RNA isolation and cDNA synthesis*

Total RNA was isolated from the liver or perfused hepatocytes via TRIzol extraction. One milliliter of TRIzol (Invitrogen0 was added to small pieces (approximately 30 mg to 50 mg) of either snap-frozen liver or hepatocyte isolate. Lysing of the cells was hastened by homogenization via bullet blending with NextAdvanced Zirconium oxide 1mm beads. Chloroform was added, and the mixture was centrifuged for 10 minutes at 10,000xg, which caused a separation of layers. The aqueous layer containing extracted RNA was removed, and to this was added 600 μL of isopropanol. The solution was mixed and stored at -20℃ overnight (or for at least 8 hours). Afterward, the mixture was centrifuged for 40 min at 12,000xg, causing the precipitation of the RNA. The chloroform was removed, and the RNA pellet was washed with 70% ethanol. The RNA was then dissolved in water. RNA purity and concentration were assessed with a Biotek Synergy 2 UV spectrometer.

cDNA synthesis was performed on 1 μg of RNA, using random hexamer primers and Maxima Reverse Transcriptase (Thermo Fisher Scientific) following manufacturer protocol. The cDNA was diluted to a total volume of 200 μL.

*RNA-seq*

RNA was isolated from hepatocytes using an RNeasy tissue mini-kit (Qiagen). Before library preparation, RNA quality was assessed using an Agilent Bioanalyzer by the Functional Genomics Core at the Roy J. Carver Biotechnology Center, UIUC. Poly-A selected, RNA-seq libraries preparation, and 150-bp paired-end Illumina sequencing were performed on a NOVASEQ 6000 at the High Throughput Sequencing and Genotyping Unit, UIUC. RNA-seq reads were processed for quality and read length filters using Trimmomatic (version 0.39). RNA-seq reads were further aligned to the mouse genome (mm10) using STAR (version 2.5.2).

Gene expression levels were determined as TPM using count and differential expression values obtained from DESeq2 (version 1.8.2), Htseq (version 0.6.1), and Cuffdiff 2 (version 2.2.1) [6–8]. Genes were considered as having significant differential expression following imposed cutoff clearance (FDR $\leq$ 0.05, log2(fold change) $\geq$1). Differential splicing analysis was performed using rMATS (version 3.2.5), and significant events were identified using imposed cutoffs (FDR $\leq$ 0.10, junction read counts ≥ 10, PSI ≥ 10%) [9]. Motif analysis for differentially spliced exons was performed using rMAPS with default parameters and putative motifs as described previously [10]. To perform alternative polyadenylation (APA) analysis, 3'UTR expression quantification was performed via Salmon (version 1.0.0), followed by analysis via qAPA (version 1.2.2) [11,12]. APA events were determined as significant if a change of 5 TPM or greater. Gene ontology analysis was performed using DAVID (version 6.8) [13–16].

To perform the exon ontology analysis, exons undergoing significant changes in splicing were converted to corresponding human exons in the hg19 annotation using UCSC liftover with a minimum ratio of bases matching 0.8[17]. Additionally, the exons in hg19 reported by UCSC liftover were checked for gene identity corresponding to the mouse exon's parent gene. These exons were then analyzed for ontology using the exon ontology and FasterDB packages [18].

*q-RT-PCR*

Relative gene expression analysis was performed with ~50 ng of cDNA per reaction using a SYBR® Green™ assay for Real-Time Quantitative Reverse Transcription PCR (qRT-PCR). cDNA, primer, SYBR, and water were mixed in a qPCR plate. The qPCR reaction was performed using a QuantStudio 3 Real-Time PCR System (Thermo Fisher). Standard cycling conditions for a SYBR® Green™ based $\Delta\Delta CT$ assay with a reduced elongation step of 35 seconds were used. β-actin was used as a loading control unless otherwise noted. Relative gene expression is listed as relative to β-actin for quantification of the DT960i Primers used for the qPCR reactions are listed in Supplemental Table 1.

*RNA-FISH and immunofluorescence (IF)*

At sacrificing, pieces of liver were placed within the Tissue-Tek OCT (Optimal Cutting Temperature) Compound and frozen with dry ice. These frozen blocks were sectioned at 10 μm with a Leica CM3050 S cryostat at the Carl R. Woese Institute for Genomic Biology (UIUC).

RNA-FISH/IF was performed on these cryosections. Sections were washed in 1X PBS and then fixed with 10% NBF for 15-30 min. Slides were washed with 1X PBS, permeabilized with 0.5% Triton-X in 1X PBS for 10 minutes, washed with 1X SSC, and then washed with 30% Formamide in 2X SSC. FISH probe was then applied (the solution contained 2 μg/mL BSA, 66 μg/mL yeast tRNA, and 1 ng/μL Cy5-$(CAG)_{10}$ (Integrated DNA Technologies) dissolved in 30% formamide in 2X SSC). After incubation at 37℃ for two hours, sections were washed with 30% formamide in 2X SSC (for 30 minutes) and washed twice with 1X SSC. The slides were again fixed with 10% NBF (for 10 min), washed with 1X TBS, and re-permeabilized with 0.5% Triton-X in PBS (again for 10 min). Slides were washed in 1X TBS before being blocked in 10% normal goat serum with 1% BSA in 1X TBS for two hours (at room temperature). After blocking, the slides were drained with a vacuum trap and incubated in 1:500 primary antibody (anti-Mbnl1 [sc-

47740] or anti-Mbnl2 [sc-136167] from Santa Cruz Biotechnology) in 1X TBS with 1% BSA at 4℃ overnight.

The next day, the slides were washed in 1X TBS with 0.05% Triton-X and then washed in 1X TBS. Then, they were incubated in 1:500 secondary antibody (anti-Mouse IgG conjugated to DyLight 488 [AB_1965946] from Thermo Fisher Scientific) in 1X TBS for 1 hour at room temperature. Slides were washed with TBS with 0.05% Triton-X, washed with 1X PBS, stained with NucBlue (Invitrogen) in 1X PBS for 20 minutes, and washed with 1X PBS. Slides were imaged on a Zeiss LSM 710 microscope at the Carl R. Woese Institute for Genomic Biology at UIUC. A list of antibodies used is provided in Supplemental Table 2.

*RT-PCR splice assays*

Target events were amplified via PCR using the primers listed in Supplemental Table 1. The PCR cycle had a melting temperature of 95℃, an annealing temperature of 55℃, and an elongation temperature of 72℃. The PCR product was resolved down a 5.5% PAGE gel, stained with ethidium bromide, and imaged using a Bio-Rad Gel Doc machine.

Bio-Rad Image Lab Software (version 6.0.1) was used to measure the intensity of the bands appearing on the gel to quantify the percent splicing change. Splicing change is reported as Percent Spliced In, a ratio of the intensity of the upper band (containing the alternatively spliced exon) to the combined intensity of the upper and lower bands (the lower band does not include the alternatively spliced exon).

*Histology: hematoxylin and eosin*

To perform hematoxylin and eosin staining, paraffin-embedded tissues were sectioned into 5 μm thick sections and then deparaffinized with three xylene washes. The slides were rehydrated in ethanol solutions of decreasing concentration (100%, 95%, 80%, and 50%) before being placed in water. The slides were stained with Hematoxylin 7211 for 1.5 to 2 minutes and washed in water. Slides were blued in a 2% sodium bicarbonate and 0.2%

magnesium sulfate bluing solution before being soaked in water again. After being placed in an ethanol solution, the slides were stained with eosin for 15-20 seconds, washed with ethanol, and dehydrated with xylene. They were finally coverslipped with Permount.

All slides were imaged on a Hamamatsu Nanozoomer at the Carl R. Woese Institute for Genomic Biology at UIUC.

##### *Histology: oil red O. staining*

Oil red O. stain was performed on cryosections (reference “RNA-FISH and Immunofluorescence (IF)” for details on the preparation of cryosections). The cryosections were first fixed in 10% NBF and then hydrated in 1X PBS. After placing the slides in 60% isopropanol, the slides were stained in fresh Oil Red O. solution (consisting of Oil Red O. dye dissolved in 60% isopropanol). The slides were then washed with 60% isopropanol, counterstained briefly in Hematoxylin 7211, and washed with tap water. The slides were then mounted with CC mount. The slides were imaged on a Hamamatsu Nanozoomer at the Carl R. Woese Institute for Genomic Biology at UIUC.

To quantitatively measure the accumulation of lipids, Oil Red O. Images at 10X zoom (approximately 1.75 x 1 mm), a minimum of 3 per sample, were collected and then analyzed by measuring the relative volume of red channel compared to blue (using standard RGB-based image splitting) using a pipeline on Cell Profiler [19,20].

##### *Serum alanine aminotransferase (ALT) and aspartate aminotransferase (AST) testing*

Whole blood from mice was collected via retro-orbital puncture in Capiject gel/clot activator tubes, centrifuged for 3 min at 3000 × *g*, and then stored at −80 °C till further analysis. ALT and AST analysis is performed using commercial Thermo Scientific serum chemistry kits. Measurements were made in duplicate.

*Zoxazolamine recovery testing*

Zoxazolamine (Zox) was purchased from Sigma-Aldrich (A45807). Zox solution was prepared the day before experimentation by dissolving enough Zox in DMSO such that the final concentration of Zox was 15 ug/µL in a 95% Corn-Oil, 5% DMSO solution. Mice then fasted for 18-22 hours before 120 mg/kg Zox injections; Zox solutions were homogenized vigorously between injections to ensure a homogenous solution.

Once treated with Zox, mice could roam freely until motor function was lost, at which point mice would be placed in the supine position on an insulating blanket. Time would then be measured until the mice regained muscle control. Time counting ceased when the mouse could successfully self-right three times[21].

*APAP insult testing*

Acetaminophen (APAP) was purchased from Sigma-Aldrich (A5000). Solutions were prepared immediately before use. Twenty mg of APAP would be dissolved in one mL of sterile 1x PBS. This solution would then be heated at 55C for 15 min with periodic vortexing. Once the 15-minute incubation was complete, the APAP solution was kept at 40C for the injection process, and vigorous mixing occurred between injections.

Mice for APAP testing were fasted overnight (18-22 hours) and then injected with 350 mg/kg. Mice were observed for 8 hours when the surviving mice were returned to the mouse facilities. The following day, mice would be harvested for serum and liver samples 24 hours post APAP injection [22,23].

*Western blot and antibodies*

Approximately 50 mg of liver tissue was homogenized via bullet blending and lysed via sonication in RIPA lysis buffer (1x SDS, 1% v/v phosphatase inhibitor cocktail 3 (Sigma-Aldrich), 1x Pierce protease inhibitor mini tablets (Thermo Scientific). Samples were then separated with SDS-PAGE and transferred overnight in Towbin buffer with 0.5% SDS. Blots were then visualized by ECL after incubating with primary and secondary antibodies. All primary antibodies were diluted in AdvanBlock-Chemi Blocking solution from Advansta, and secondary antibodies were diluted in TBST (0.1% v/v Tween-20). See supplementary table 4 for a list of antibodies and suppliers.

*Lipid extraction and analysis*

Twenty to fifty milligrams of liver tissue were weighed into locking cap 1.5 mL Eppendorf tubes. To this was added 600 μL chilled 3:2 hexanes/isopropanol solution and 6-10 NextAdvanced zirconium oxide 1mm beads. The tubes were homogenized on a bullet blender for 30 seconds and then allowed to rest on ice for 2 minutes. This homogenization process was repeated five times before samples were spun down at *3000 x g* for 10 minutes at 4℃. The liquid phase was then collected into a second Eppendorf tube and set aside. An additional 600 μL chilled 3:2 hexanes/isopropanol solution was then used to break up the pellet, and samples were rested on ice for 15 min with periodic vortexing. Again, samples were centrifuged at *3000 x g* for 10 min, and the liquid phase was then combined with the liquid set aside previously. Samples were then allowed to dry in the open air overnight.

Samples were diluted in 200 μL of 1x PBS with 2% Triton X-100. Samples were vortexed and allowed to dissolve at 4C; further dilution would occur if necessary. Once dissolved, samples were analyzed using an Infinity triglycerides colorimetric kit.

*Statistical analysis*

All quantitative experiments have at least three independent biological repeats. The results were expressed with mean and standard deviation unless mentioned otherwise. All western blots and qPCR experiments were performed at least three times per sample, and all serum tests, GTT, and ALT/AST tests were performed at least twice per animal and averaged. Differences between groups were examined for statistical significance using unpaired T-tests when comparing directly between groups or one-way analysis of variance (ANOVA) for more than two groups using the GraphPad Prism 9 Software. Statistical outliers were determined with the ROUT method in Prism, with Q = 5%. P-value $< 0.05$ or FDR $< 0.10$ was considered significant. In all figures, significance was set as $p < 0.05$, "*" indicates $p < 0.05$, "**" indicates $p < 0.01$, "***" indicates $p < 0.001$, and "****" indicates $p < 0.0001$.
